# A living conductive marine biofilm engineered to sense and respond to small molecules

**DOI:** 10.1101/2022.08.23.504934

**Authors:** Lina J. Bird, Dasha Leary, Judson Hervey, Jaimee Compton, Daniel Phillips, Leonard M. Tender, Christopher A. Voigt, Sarah M. Glaven

## Abstract

Engineered electroactive bacteria have potential applications ranging from sensing to biosynthesis. In order to advance the use of engineered electroactive bacteria, it is important to demonstrate functional expression of electron transfer modules in chassis adapted to operationally relevant conditions, such as non-freshwater environments. Here, we use the *Shewanella oneidensis* electron transfer pathway to induce current production in a marine bacterium, *Marinobacter atlanticus*, during biofilm growth in artificial seawater. Genetically encoded sensors optimized for use in *E. coli* were used to control protein expression in planktonic and biofilm attached cells. Significant current production required addition of menaquinone, which *M. atlanticus* does not produce, for electron transfer from the inner membrane to the expressed electron transfer pathway. Current through the *S. oneidensis* pathway in *M. atlanticus* was observed when inducing molecules were present during biofilm formation. Electron transfer was also reversible, indicating electron transfer into *M. atlanticus* could be controlled. These results show that an operationally relevant marine bacterium can be genetically engineered for environmental sensing and response using an electrical signal.

## Introduction

Microbial biofilms that naturally catalyze electron transfer at electrodes have been used for applications such as energy harvesting by microbial fuel cells ^1-3^ and reduction of carbon dioxide (CO_2_) during electrotrophy ^4-5^. Efforts to place natural extracellular electron transfer (EET) pathways under genetic control to act as electrical reporters when expression is activated by chemical inducers have been demonstrated in both the native host species^6-7^ and in *Escherichia coli* ^8-11^. For expression of the EET pathway in *E. coli*, protein expression is initiated during the pre-growth phase in rich medium, and introduced into the reactor at a higher cell density ^9^. Progress has been made towards inducing a fast response of the resulting circuit for electrical reporting; however, cells require extensive engineering and encapsulation to be located at the electrode surface ^12^. In addition, *E. coli*, like *S. oneidensis*, may not function optimally in certain environments, particularly in marine and high saline conditions.

*Marinobacter atlanticus* CP1 is a biofilm-forming, marine bacterium isolated from an electroactive microbial community enriched from the Atlantic Ocean ^13^. *M. atlanticus* naturally produces small amounts of electrical current from oxidation of organic acids ^14-15^ when grown in an electrochemical reactor even in the presence of O_2_. The mechanism for this native EET is not fully understood, and *M. atlanticus* does not contain large, multi-heme *c*-type cytochromes found in *S. oneidensis* and other high performing electricigens. Extensive electrochemical ^14^ and transcriptomics ^16^ analysis suggests that EET in this organism is facilitated in part by diffusion of redox-active species between the cell and electrode. The low intrinsic current density, lack of native multi-heme *c-*type cytochromes, its ability to form robust biofilms, and available genetic system ^17^ make *M. atlanticus* suitable for development as a living electronic sensor for applications in the marine environment ^18^.

In this study, we transferred the *So*EET pathway to *M. atlanticus* as a step toward a living electrical reporter system for the marine environment. Using a suite of genetically encoded sensors originally optimized for *E. coli*, we demonstrated functionality across a range of inducer concentrations for activation of expression of *So*EET proteins (MtrA, MtrB, MtrC, CymA, and CctA) both from the genome and from plasmids. Protein expression, and subsequent current generation, occurred when inducers were added at the time of inoculation or (with a less robust response) after 24 hours following electrode biofilm formation. Expression of the *So*EET pathway resulted in reversible electron transfer in *M. atlanticus* with O_2_ as the terminal electron acceptor; this represents an important step towards bidirectional electron transfer. A key factor in engineered EET was the addition of menaquinone-4 (MK-4), a required cofactor for electron transfer through CymA. The results presented here demonstrate a novel marine chassis for operationalizing synthetic biology in the marine environment and provide new design rules for engineering electroactive bacteria.

## Results

### Tunable, genetically encoded sensors for *Marinobacter atlanticus* CP1

We evaluated activation and dynamic response of seven different genetically encoded sensors in *M. atlanticus* to be used to control expression of genes for the *So*EET pathway (Figure 1A). These sensors had previously been evolved for high dynamic range and low cross-reactivity in *E. coli* ^19^, and were not further optimized prior to testing in *M. atlanticus*. Sensors were cloned into a broad host range plasmid (pBBr) for conjugative transfer into the wild type *M. atlanticus* (CP1-wt) background strain yielding strains that allow the independent control of gene expression by applying 2,4-diacetylphophloroglucinol (DAPG), cuminic acid (Cuma), vanillic acid (Van), isopropyl β -d-1-thiogalactopyranoside (IPTG), tetracycline (Tc), choline chloride (Cho), or naringenin (Nar) to the growth media. Each sensor consists of a weak constitutive promoter (P_LacI_) driving the expression of the regulatory gene and an output promoter that is acted on by the regulator. Sensors are activated when a specific inducer molecule binds to the regulator releasing it from the promoter-binding site and in this case, initiating expression of yellow fluorescent protein (YFP).

**Figure 1.**
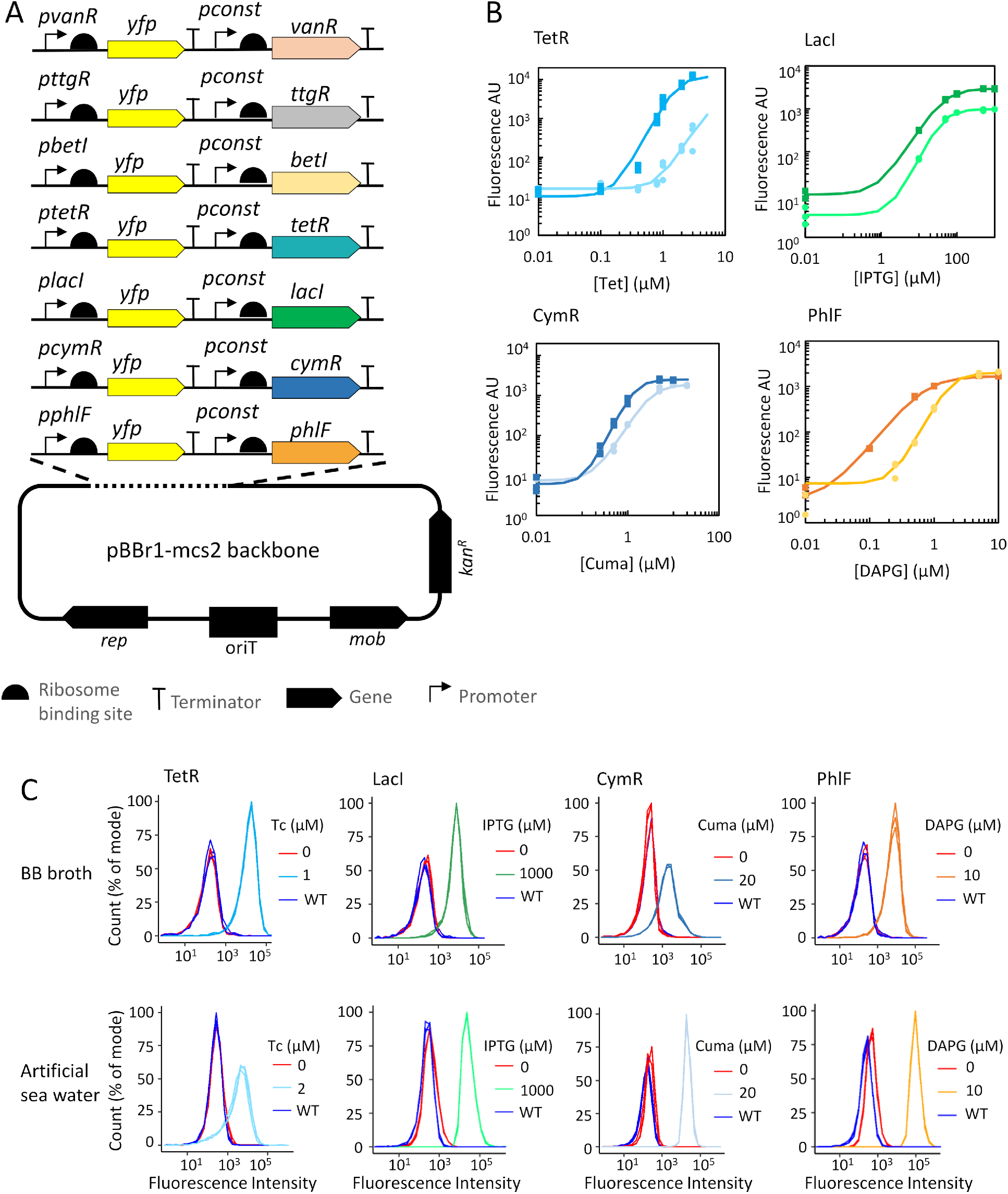
Evaluation of sensors from the Marionette promoter library in *Marinobacter atlanticus*. A) Genetic configuration of sensors in a broad range host vector (pBBr). B) Response function of Tc (light blue), IPTG (green), Cuma (blue), and DAPG (orange) sensors at 21 hours post inoculation in either rich (BB broth, darker shade) or minimal (low-Ca artificial seawater, lighter shade) medium when activated during exponential growth phase (6 hours) across a range of inducer concentrations. Parameters of the fitted binding curves are reported in Supplemental Table 2. C) Cytometry histograms for the Tc, IPTG, Cuma, and DAPG sensors comparing uninduced and maximum induction to wild type *M. atlanticus* in rich (BB broth) and minimal (low calcium artificial sea water) medium. For each condition, n=3, and replicates are plotted separately.

Sensor activity in the absence of inducer (leakiness) and response function were initially evaluated during exponential growth in both rich (BB medium) and minimal (artificial seawater (ASW)) media by plate reader assay. Inducer molecules were added 6 hours post-inoculation and fluorescence intensity is reported here at 21 hours post-inoculation (Figure 1B). As indicated by the parameters in Supplemental Table 2, sensors for DAPG, Cuma, IPTG, and Tc showed low background expression of YFP and a dynamic range of 2-3 orders of magnitude, with a change in inducer concentration in both types of medium. The dynamic ranges were generally slightly lower than those seen in *E. coli* with the Marionette system, while the K-values, and hence the concentrations at which the small molecules could be detected, was comparable to that seen in *E. coli* ^19^. The Nar sensor showed a response in rich medium (Supplemental Figure 1A), but with a limited dynamic range due to cellular toxicity (Supplemental Figure 1B) and was not tested further. Sensors for Van and Cho were also tested but showed little response (Supplemental Figure 1A) and were also not evaluated further.

Flow cytometry was used to evaluate sensor activation across the entire cell population for DAPG, Cuma, IPTG, and Tc sensing strains at the concentration resulting in maximum fluorescence in plate reader assays (Figure 1C). The majority of cells from the DAPG, IPTG, and Tc responsive strains were induced in rich medium; however, the Cuma sensing strain displayed a broader distribution of fluorescence intensity with some portion of cells remaining uninduced. In minimal medium, the induced Cuma sensing strain had clear separation from the uninduced cells while the Tc sensing strain had a broader distribution. Histograms for the uninduced DAPG sensing strain relative to wild type *M. atlanticus* indicated potential promoter leakiness as the peak fluorescence was shifted slightly.

Sensor activation was also evaluated in liquid cultures during stationary phase (induction after 18 hours growth), resulting in an order of magnitude lower YFP intensity compared to log phase for the IPTG, Cuma, and DAPG sensors, and little to no response for the Tc sensor (Supplemental Figure 2, Supplemental Table 2). This result suggested that sensor activation during biofilm growth, when sensors must be used to activate expression of genes for EET proteins, may be dramatically different than during planktonic, log-phase growth.

Based on both the dynamic response and inducibility of the cell population in artificial seawater, we selected the IPTG and DAPG sensors for inducible biofilm expression of the genes for the *So*EET proteins in *M. atlanticus*.

### *Cloning and expression of the* S. oneidensis *EET proteins*

Current production through the *So*EET proteins in *E. coli* requires expression of the outer membrane metal-reducing (*mtr*) operon ^20-21^, MtrCAB, the small tetraheme *c-*type cytochrome CctA ^22^, the inner membrane *c*-type cytochrome CymA ^23-24^, and a copy of *E. coli’s* native cytochrome *c* maturation (*ccm*) system under a constitutive promoter ^25^. We sought to determine whether the same components were necessary in *M. atlanticus*.

First, we inserted the genes for MtrCAB into genome position 1926204, a neutral site in the *M. atlanticus* genome, under control of the IPTG sensor (CP1-mtr). Next, we constructed a vector using the broad host range plasmid pBBr for expression of *cymA* and *cctA* under control of the DAPG sensor and transferred it into the Mtr background strain (CP1-mcc). Finally, we constructed a second vector containing a copy of the native *M. atlanticus ccm* operon under control of the Tc sensor using the broad host range vector pSEVA 351 ^26^, and transferred this into the CP1-mcc strain to create CP1-eet. A summary of the constructs and their role in EET is depicted in Figure 2A and B, and the strain names and genetic alterations are summarized in Table 1.

**Table 1:** strains and plasmids used in this study.

| Strain | Description | Source |
| --- | --- | --- |
| CP1 | *M. atlanticus* CP1 wild type strain | 17 |
| CP1:dTomato | CP1 with constitutively expressed dTomato fluorescent protein inserted at the neutral site | 17 |
| CP1:MtrCAB | CP1 with the MtrCAB operon under the control of the LacI expression cassette, inserted at the neutral site. | This study |
| *E. coli* DH5a | Plasmid maintenance strain of *E. coli* | 58 |
| *E. coli* WM3064 | Conjugation strain of *e. coli*; DAP auxotroph | 59 |
| **Abbreviated strain names in electrochemical experiments** |  |  |
| Cp1-mcc | CP1:MtrCAB + pLB7 (the pBBR plasmid containing *cymA* and *cctA* under control of the DAPG sensor) | This study |
| CP1-eet | CP1:Mtr+pLB7 with the pSEVA351 plasmid containing the *ccm* operon under control of the aTc sensor | This study |
| CP1-yfp | *M. atlanticus* + pLB35 (phlF-yfp) and pLB43 (*ccm* pathway) | This study |
| CP1-mtr | CP1:MtrCAB with pLB43 (*ccm* pathway) | This Study |
| CP1-cymcct | *M. atlanticus* CP1 with pLB7 (*cymA* and *cctA* with phlF promoter) and pLB 43 (*ccm* pathway) | This Study |
| CP1-ccm | *M. atlanticus* with pLB42 (*ccm* pathway) | This Study |
| CP1-wt | *M. atlanticus* CP1 wild type strain | 17 |
| **Plasmid** | **Description** | **Source** |
| pBBR1mcs2 | Broad range plasmid backbone with kanamycin resistance | 60 |
| pSEVA351 | Broad range vector with chloramphenicol resistance | 26 |
| pSEVA651 | Broad range vector with gentamycin resistance. | 26 |
| pJQ200sk | Narrow range suicide vector used to create genome integrations. Gentamycin selection and sacB counter selection. | 61 |
| pLB33 | pBBr1mcs2 backbone with TetR-yfp cassette | This study |
| pLB36 | pBBr1mcs2 backbone with LacI-yfp cassette | This study |
| pLB35 | pBBr1mcs2 backbone with PhlF-yfp cassette | This study |
| pLB34 | pBBr1mcs2 backbone with CymR-yfp cassette | This study |
| pLB37 | pBBr1mcs2 backbone with TtgR-yfp cassette | This study |
| pLB45 | pBBr1mcs2 backbone with BetI-yfp cassette | This study |
| pLB7 | *cymA and cctA* under PhlF control | This study |
| pLB9 | *cymA-cctA* operon under PhlF control | This study |
| pLB19 | mtrCAB operon under control of LacI on pJQ200sk backbone for genome integration | This study |
| pLB43 | *M. atlanticus* cytochrome *c* maturation pathway under control of tetR promoter on pSEVA651 backbone with gentamycin resistance | This study |

**Figure 2.**
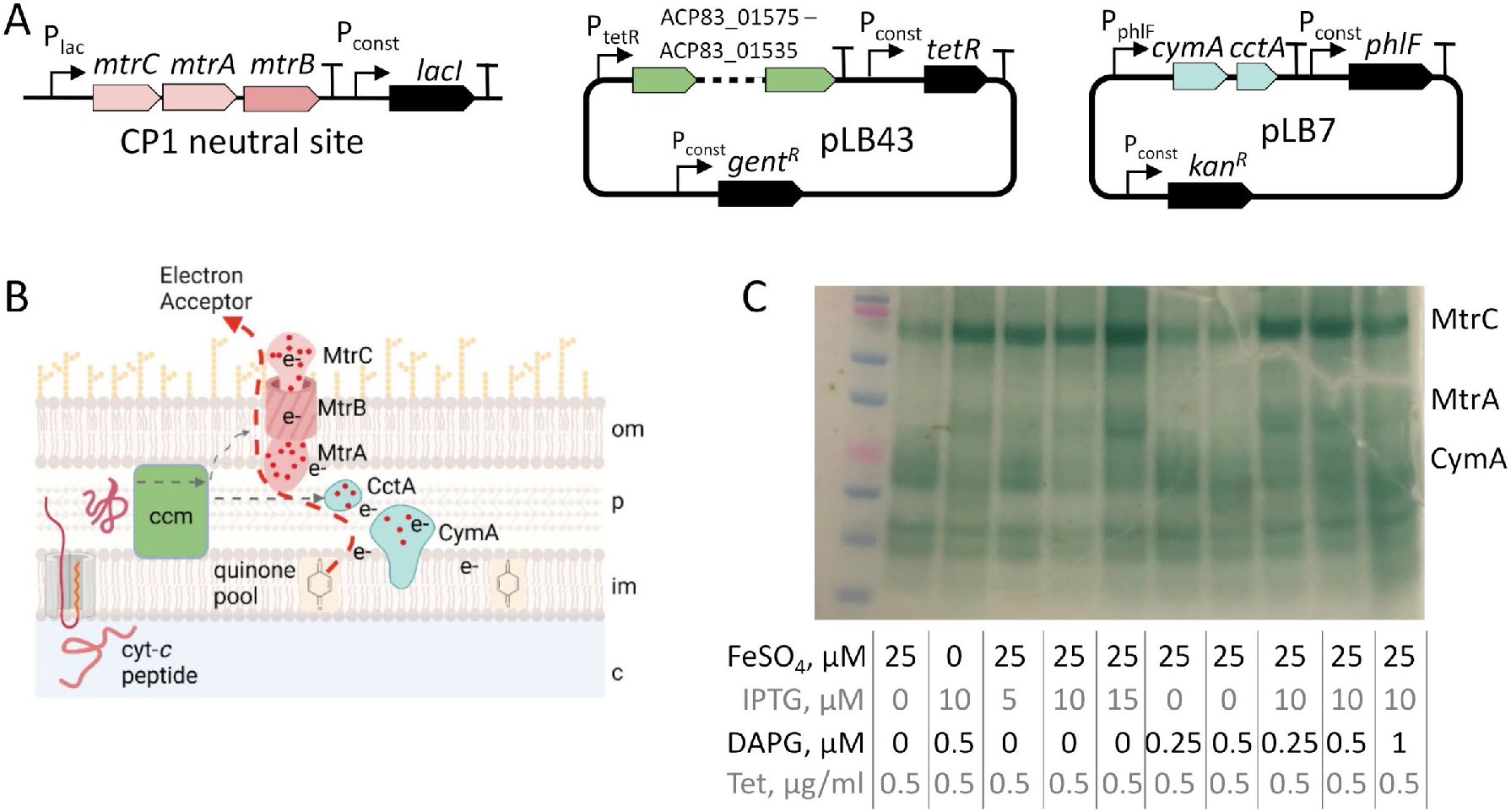
Construction and evaluation of electron transfer proteins from *S. oneidensis* in *M. atlanticus*. **A)** Design of strain CP1-eet for expression of extracellular electron transfer proteins from *S. oneidensis* in *M. atlanticus*. The *mtr* operon was expressed from a neutral site on the *M. atlanticus* genome under control of the IPTG sensor. *cymA* and *cctA* were expressed from a plasmid under control of the DAPG sensor, and the cytochrome *c* maturation pathway was expressed from a complimentary plasmid under control of the Tc sensor. **B)** Schematic of the *So*EET pathway components based on their function in *S. oneidensis*. Electrons travel from the quinone pool through CymA to CctA (light blue), and are then transferred to the MtrCAB complex (pink), in which the cytochromes MtrA and MtrC are connected by the outer membrane porin MtrC. The cytochrome *c* maturation (Ccm) system (green) is required to correctly process the cytochromes in the pathway. **C)** Expression of cyt-*c* proteins was evaluated by heme staining following gel electrophoresis.

DAPG, IPTG, and Tc were added to the growth medium at the lowest concentrations required to detect YFP fluorescence to test for expression of *So*EET proteins. As has previously been noted for *E. coli*, cell pellets from cultures expressing both the MtrCAB complex and the additional copy of the *ccm* operon showed a deeper red hue than un-induced pellets due to the increase in heme-containing *c-*type cytochrome proteins (Supplemental Figure 3A). Expression of CymA and CctA did not result in visibly redder pellets. Protein expression was evaluated after growth in artificial seawater medium by heme stain following gel electrophoresis for CP1-eet under varying inducer concentrations (Figure 2C). When increasing concentrations of IPTG were added, band intensity for *c-*type cytochrome proteins with similar molecular weights to MtrC and MtrA increased. This pattern was the same in rich medium (Supplemental Figure 3B). Despite the tight regulation of YFP under the *lacI* promoter, bands for MtrC and MtrA were detected even when IPTG was not added to the medium. A band with similar molecular weight to CymA was visible only when DAPG was added to the medium. CctA, the smallest of the 4 cytochromes, was not detected. Induction of either sensor with higher concentrations of inducer led to growth inhibition (not shown). Addition of iron sulfate (FeSO_4_) had a slight effect on band intensity. However, the heme stains used here were not quantitative, and these patterns would need to be evaluated further before tunability could be confirmed. Protein expression was confirmed following growth and induction in rich medium for MtrC, MtrB, and CymA by high performance liquid chromatography with mass spectrometry (HPLC-MS) (Supplemental Figure 3C).

### Sensor activation in electrode biofilms

Next, we tested whether the DAPG, Tc, and IPTG sensors could be activated in electrode biofilms, rather than planktonic cells. YFP expression was tested in a background strain of *M. atlanticus* with the red fluorescent protein (dTomato) inserted into the neutral site in order to visualize all biofilm cells. Inducers (10 µM and 1 mM IPTG, 0.25 and 10 µM DAPG, 0.5 and 1 µg/mL Tc) were added to standard bioelectrochemical reactors (schematic in Supplemental Figure 4) at the time of inoculation under current-producing conditions (0.510 V versus standard hydrogen electrode (SHE)). Sensor activation was evaluated after 24 hours of biofilm growth by removing the electrode from the reactor and performing laser scanning confocal microscopy (Figure 3). Expression of YFP was detected in the presence of the inducer but not when the inducer was absent. YFP expression was observed in fewer cells in the Tc responsive strain, suggesting that the TetR promoter may not be as robust during biofilm growth. Overall, results show that sensor activation is possible even when cells grow attached to an electrode surface.

**Figure 3.**
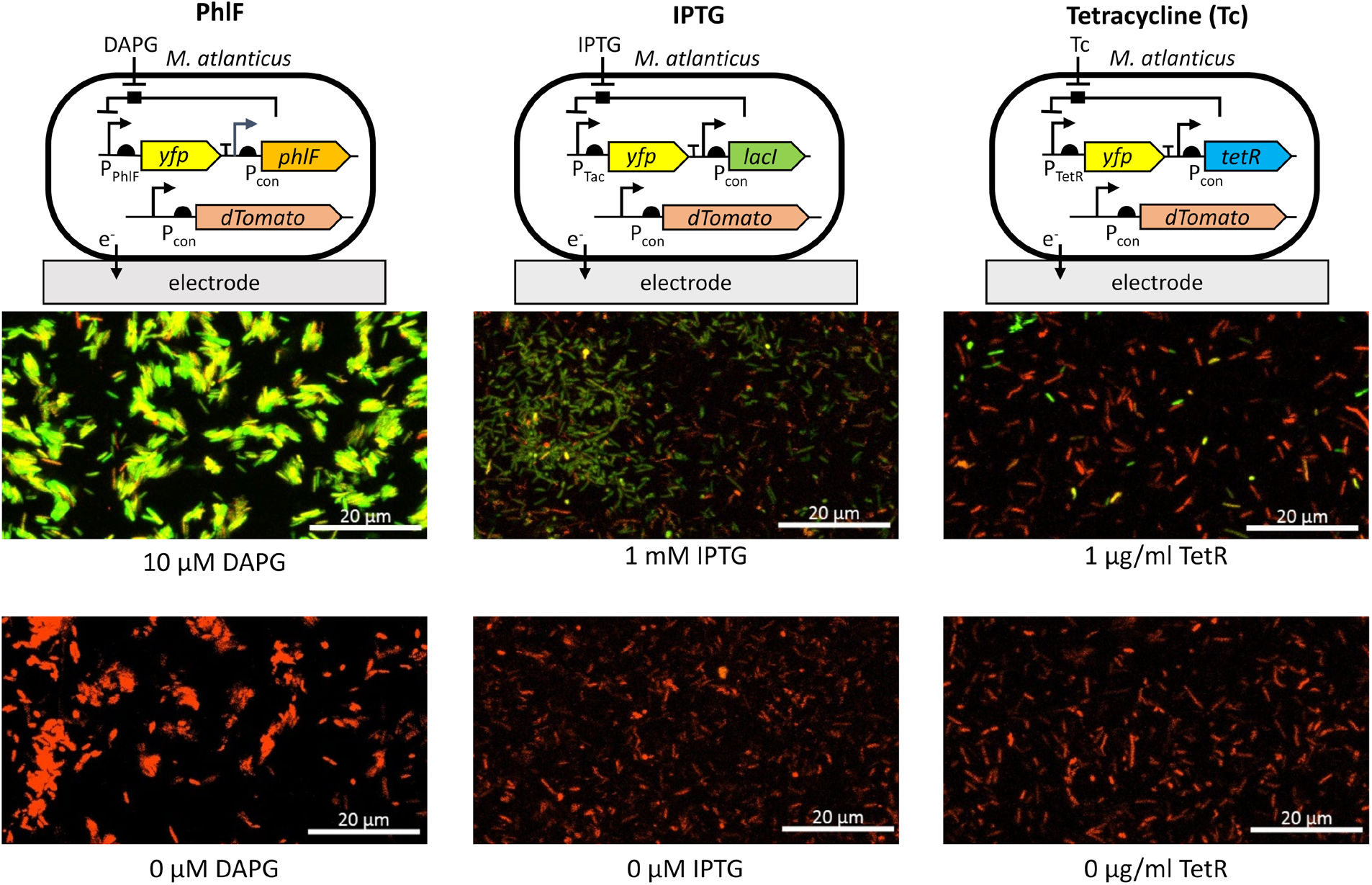
Activation of genetically encoded sensors during biofilm growth in a standard bioelectrochemical reactor. Biofilms were formed on carbon cloth working electrodes (20 cm^2^) with an applied potential of 0.510 V vs. the standard hydrogen electrode (SHE) and were imaged using confocal microscopy 24 hours after inoculation. For each strain, the indicated inducer was added at the time of inoculation at the following concentrations: 10 µM and 1 mM IPTG, 0.25 µM and 10 µM DAPG, 0.5 µg/mL and 1 µg/mL Tc. The *M. atlanticus* background strain contained a genomic insertion of the gene for the red fluorescent protein dTomato. Red channel = dTomato, green channel = YFP. Yellow color is the overlay of the two channels. Images are maximum intensity projections, representative of 2 independent experiments for each strain.

### Current production in M. atlanticus engineered to express the SoEET proteins

Next we tested whether expression of the *So*EET proteins and extra copy of the *ccm* pathway results in increased current in *M. atlanticus*. Sensors were activated in overnight cultures prior to inoculation into bioelectrochemical reactors. The average maximum current produced by induced CP1-eet after 50 hours was higher than CP1-wt and uninduced control (Figure 4A); however, this difference was not significant (p= 0.1 for both comparisons using the Wilcox and Gibbs test). At the end of the experiment, cyclic voltammetry (CV) was used to assess the dependency of current on the electrode potential and to characterize any associated redox features (Figure 4B). CV indicated the presence of a catalytic wave, consistent with biofilm EET ^27-28^, with an average midpoint potential (E_M_) of -0.081 V, which is in the range of that reported for MtrC by protein film voltammetry (PFV) ^29^, as well as for the MtrCAB complex measured by PFV and spectral titration in liposomes ^30^. A second feature with an average E_M_ of -0.205 V was also evident in all strains and has been observed previously for WT *M. altanticus* electrode biofilms ^14^; therefore, it is unrelated to the *So*EET proteins.

**Figure 4.**
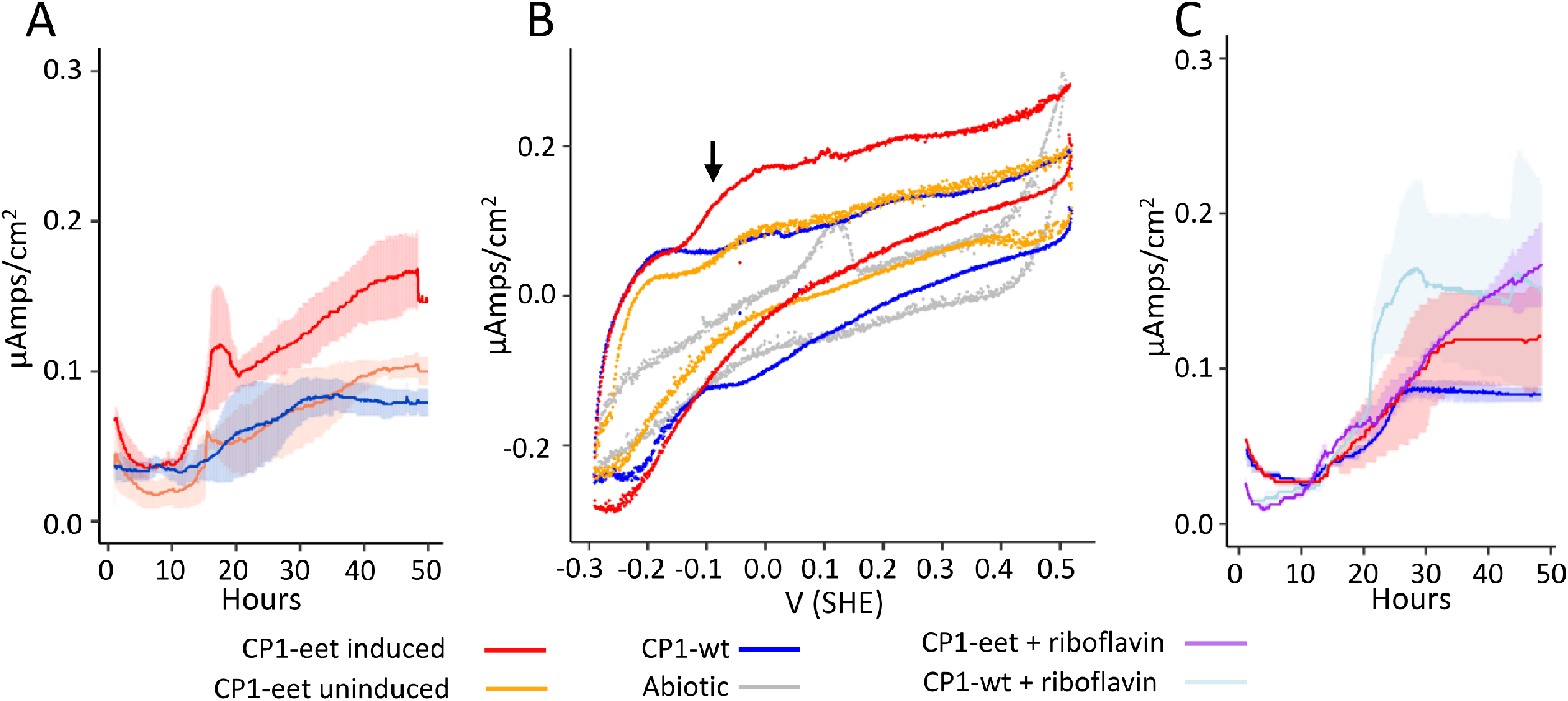
Chronoamperometry and cyclic voltammetry of engineered *M. atlanticus*. A) Average current over time of CP1-eet when sensors are activated overnight prior to inoculation (n=3) or left uninduced (n=3) compared to CP1-wt (n=3). B) Average current during cyclic voltammetry (0.2 mV/sec) performed on the reactors in panel A showing a catalytic wave (indicated by the black arrow) in the induced CP1-eet compared to controls. C) Average current over time of induced CP1-eet and CP1-yfp with and without 2 µM riboflavin added at inoculation. Shading indicates standard deviation, with n=2 for all conditions. Sensors were activated with the following concentration of inducers: 10 µM IPTG, 0.25 µM DAPG, 0.5 µg/mL Tc. Cells are added to reactors at a 1:170 dilution.

It is known that riboflavin is also required for current production in *S. oneidensis* ^31^, and increases current production in *E. coli* ^10^. However, addition of riboflavin to our reactors did not increase the current in CP1-eet (Figure 4C).

### Menaquinone is required for higher current

Since the EET proteins appear to be present in CP1-eet, shown both by the protein analysis and CV described above, we hypothesized that an additional factor was limiting current. In *S. oneidensis*, electrons enter the respiratory chain of the inner membrane through different dehydrogenases ^32-33^ and in the absence of oxygen interact with the quinone pool, specifically menaquinone-7 ^34^. The *M. atlanticus* genome lacks key genes for the complete menaquinone pathway and instead encodes the ubiquinone biosynthesis pathway as the dominant quinone. We originally hypothesized that as O_2_ is depleted in the reactor ^17^ the equilibrium potential of ubiquinone (UQ, E_M_ 0.066 V SHE) ^35^ would shift negative due to reducing conditions at the inner the membrane, enabling ET to CymA (E_M_ -0.200 V SHE at pH 6 ^29^). However, the results presented in Figure 4 suggested electron transfer from UQ to CymA might be very limited.

Previous studies have shown that addition of exogenous menaquinone-4 (MK-4) to *S. oneidensis* rescued respiration in a mutant strain deficient in MK biosynthesis ^36^. Here, addition of MK-4 to bioelectrochemical reactors at the time of inoculation resulted in 6-fold (0.6 µA/cm^2^ vs. 0.1 µA/cm^2^) higher maximum current by the CP1-eet strain when *So*EET proteins were induced prior to inoculation (Figure 5A). This increase was significantly higher than all other conditions (p=.009 for induced vs. wild type, and .0001 for induced vs. uninduced, Wilcox and Gibbs test). Addition of MK-4 to the WT strain had no effect on current (Figure 5A, inset). We also wished to test whether all the modules we had created (*mtrCAB, cymA-cctA*, and *ccm* pathway) were necessary to see a current increase. Our tests showed no significant increase in current when CP1-mtr, CP1-cymcct, and CP1-mcc were tested with MK-4 (Figure 5A, inset); correct protein expression for these strains was confirmed via heme staining, shown in Supplemental Figure 5. Current was also 2-fold higher in the CP1-eet strain when no inducer was added (compared to wild type) and the difference was significant (p=0.03, Wilcox and Gibbs test), consistent with heme staining and further indicating that some leaky expression was occurring.

**Figure 5.**
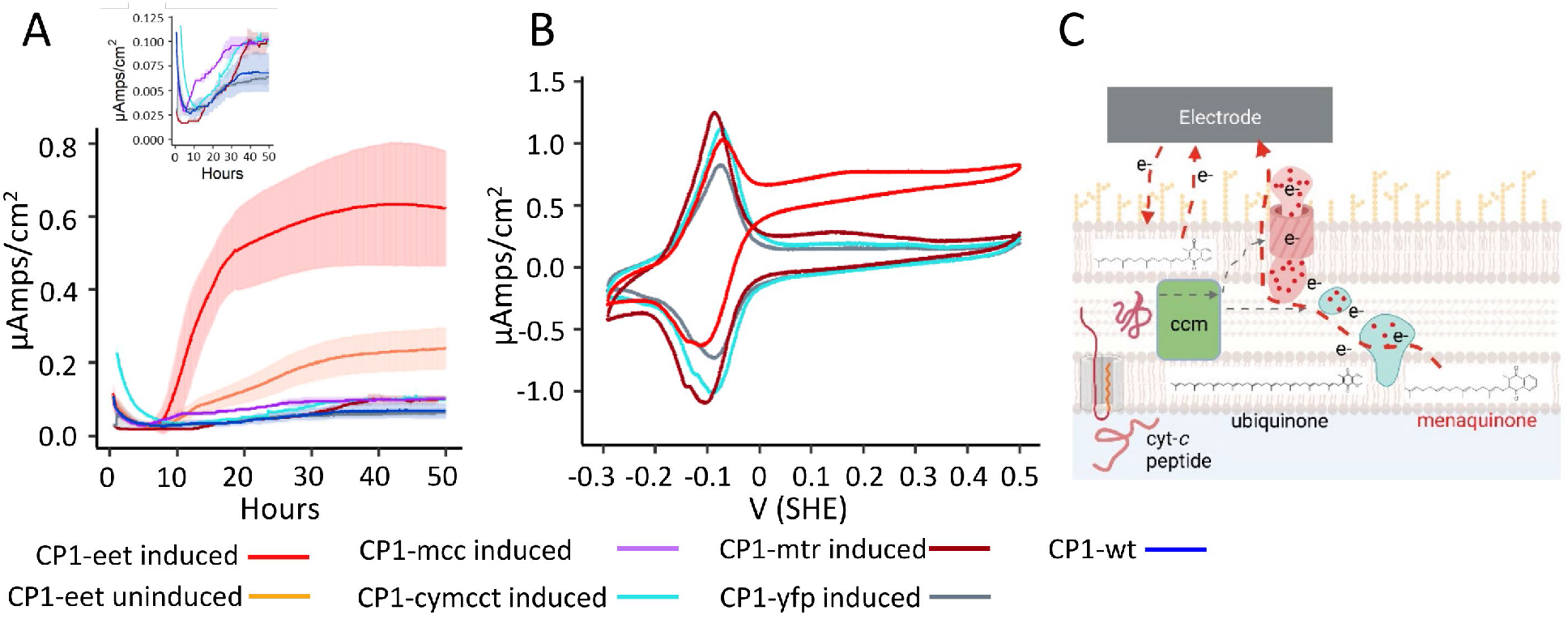
Chronoamperometry (A) and cyclic voltammetry (B) of reactors inoculated with pre-induced cultures of *M. atlanticus*. CP1-eet induced (10 µM IPTG, 0.5 µM tetracycline, 0.25 µM DAPG), n=9; CP1-eet uninduced, n=6; aqua = CP1-cymcct induced, n=2; CP1-mtr induced, n=2; CP1-yfp induced, n=3; CP1-mcc induced, n=2; CP1-wt, n=3. Shading indicates standard deviation. Inset: zoom in of low current strains. B) Representative voltammograms of CVs shown in panel A. C) Schematic depiction of MK-4 partitioning in the cell envelope within the CP1-eet strain.

Cyclic voltammetry showed a more pronounced electrocatalytic wave than in CP1-eet only (Figure 5B), consistent with the increased current generation. The addition of MK-4 to the reactors led to the appearance of a redox peak centered at -0.092 V SHE. We attribute this peak to MK-4 embedded in the cell outer membrane interacting directly with the electrode (Figure 5C). This result is consistent with previous electrochemical studies using liposomes where MK-4 was embedded and adsorbed to the electrode surface ^37^. In the absence of a biofilm, the MK-4 redox peak was shifted negative (also consistent with liposome studies) and the E_M_ was dependent on the pH of the medium (Supplemental Figure 6).

To further resolve the catalytic wave associated with the *So*EET proteins, we grew larger volumes of pre-cultures and added MK-4 to them for 2 hours before inoculating (50 mL, rinsed and concentrated) into reactors without additional MK-4. Under these conditions the MK-4 peak was greatly diminished, and the catalytic wave associated with the *So*EET proteins could be further resolved (Supplemental Figure 7). A second catalytic wave (ca. E_M_ 0.150 V SHE) associated with the *M. atlanticus* WT strain was also apparent regardless of whether MK-4 had been added to the pre-cultures. Although these high cell density cultures helped resolve catalytic features, maximum current density was not higher than typical inoculum volumes, indicating that the number of cells in the reactor was not the limiting factor in current generation.

We also tested the effect of activating sensors for different parts of the *So*EET pathway separately and over a range of concentrations in the CP1-eet strain. Addition of IPTG only (Figure 6A and 6B) induced a small amount of current, although the response was highly variable and not concentration dependent. A slightly higher concentration of IPTG led to almost no current being produced, likely due to growth inhibition by protein overproduction. Adding DAPG separately (Figure 6C and 6D) led to increased current in a dose dependent manner, although much higher variability was observed between replicate reactors than when all inducers were added. These experiments indicate that tunability of the system is possible, but requires optimization. Activation of the Tc sensor was not necessary for increased current production; however, the presence of plasmid encoded copy of the *ccm* operon was required (as shown in Figure 5A and Supplemental Figure 3). These results indicate that leaky expression of the Ccm proteins from the Tc sensor was sufficient to aid in processing the additional cytochromes *c*.

**Figure 6.**
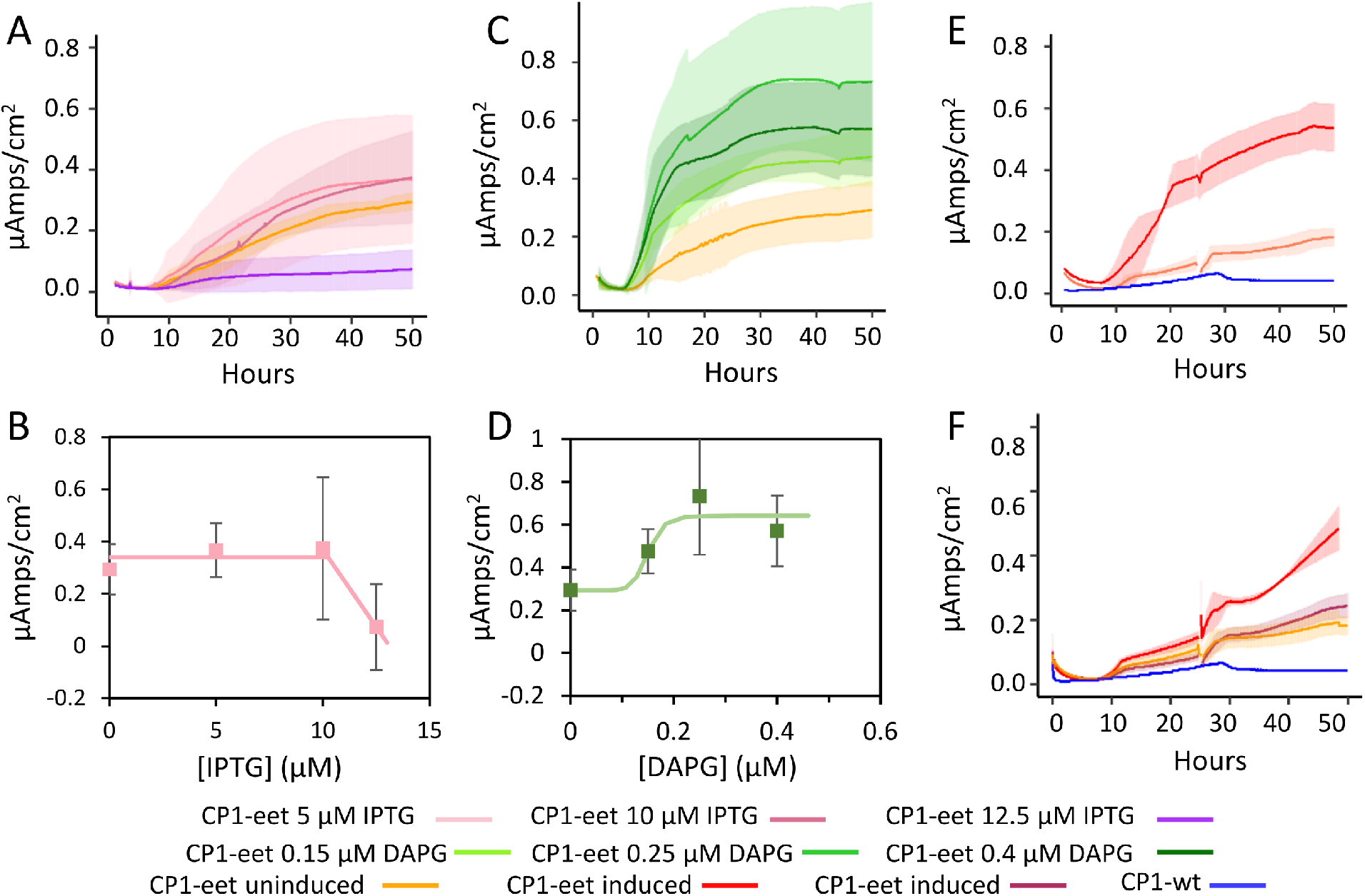
A) Sensor activation in CP1-eet with titrations of IPTG only at 5 (n=6), 10 (n=6), and 12.5 µM (n=6). B) Dose response curve of IPTG only induction. Error bars represent standard deviation of 6 replicates. C) CP1-eet titrations of DAPG only at 0.15 (n=6), 0.25 (n=6), and .4 µM (n=6). Tc was not added to any of the conditions. D) Dose response curve of DAPG titration. Error bars represent standard deviation of 6 replicates. E) Sensor activation at the time of inoculation produced currents similar to the pre-induced cultures shown in Figure 5. Induced at inoculation, n=2; uninduced, n=2; CP1-wt, n=1. Shading indicates standard deviation. F) Sensor activation and medium replacement after 24 hours in two independent experiments. Red = CP-eet induced experiment 1, n=2; maroon = CP1-eet induced experiment 2, n= 4; orange = CP1-eet uninduced with medium replaced, n=6; blue = CP1-wt (n=1).

### Current is inducible during early biofilm formation

In the electrochemical experiments described above, sensors were first activated in overnight cultures prior to inoculation into bioelectrochemical reactors. Thus, cells introduced to the reactor were primed for EET when coming into contact with the electrode. As also demonstrated above, sensors can be activated and used to express YFP in *M. atlanticus* during biofilm formation. We therefore tested whether current increased when sensors were activated during and after biofilm formation. When inducers were added to the reactor at the time of inoculation, maximum current was similar to that seen when cultures were pre-induced (Figure 6E), indicating current is not dependent on expression of the EET conduit in founder cells. When inducers were added with medium replacement once the biofilm was already formed (24 hours), as would be required for a persistent sense/respond system in the environment, the resulting increase in current was inconsistent (Figure 6F), indicating that induction of later stage biofilms is not reliable.

### Electron transfer is reversible through the Mtr pathway in M. atlanticus

In *S. oneidensis* and *E. coli*, electron transfer through the Mtr pathway is reversible ^11, 38-39^, where O_2_ or fumarate (anaerobic) serve as the terminal electron acceptor. We therefore tested whether CP1-eet had enhanced O_2_ reduction compared to the WT strain. Reactors were set up and grown as described above with CP1-wt and CP1-eet. After 24 hours, we replaced the medium in the reactors with fresh medium without lactate. A needle with a 0.2 µm filter was inserted in one of the stoppers to allow oxygen to exchange freely with the medium. CV showed that cathodic current increases at a significantly higher potential (p<0.05) in the CP1-eet strain regardless of whether sensors were activated (Figure 7A), indicating that the Mtr pathway can work in reverse, accepting electrons from the electrode. Reactors were grown both with and without MK-4. No difference was observed between reactors where MK-4 was added and those where it was not: we therefore averaged current from CVs across all reactors. The shift in onset potential for cathodic current produces was significant (p<0.05 using t-test) for applied potentials between 0.051 and 0.215 V. Once through the Mtr complex, electrons could either travel in reverse through CctA and CymA, or may bypass these proteins as they enter the quinone pool in the inner membrane, possibly through other native redox carriers (Figure 7B).

**Figure 7:**
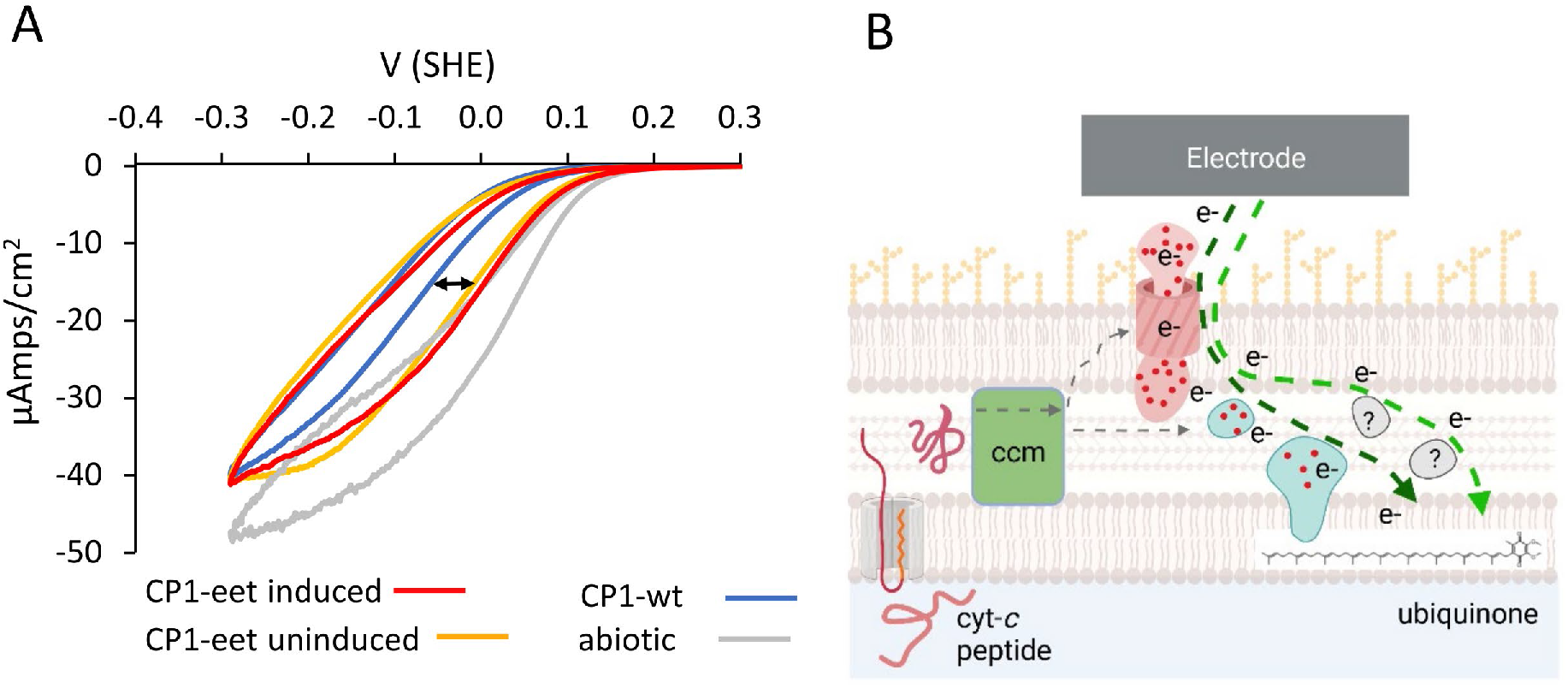
A) Average current from cyclic voltammetry performed on reactors with fresh aerobic medium added without lactate. Arrow indicates the shift in potential of negative current production in CP1-eet compared to CP1-wt. B) Model of electron transfer showing potential reverse electron flow pathways, either through the *So*EET pathway (dark green arrow) or through an unknown native pathway (light green arrow).

## Discussion

In this work, we showed that genetically encoded sensors could be deployed in the recently isolated marine chassis strain *M. atlanticus* ^13, 17^, and used to drive expression of genes for electrical current generation by direct electron transfer in seawater. Building on previous demonstrations in *E. coli*, we determined key requirements for functional expression of the *So*EET proteins, including the requirement for MK as a redox cofactor and requirement for an additional copy of the *ccm* operon. Our results demonstrate an advancement over previous efforts to refactor EET proteins from *S. oneidensis* in *E. coli* by showing expression in a marine relevant species that naturally forms electrode biofilms, and demonstrating inducibility of protein expression and increased electrical current in artificial seawater during and after biofilm formation.

IPTG, DAPG, Tc, and Cuma sensors used to induce YFP expression in both planktonic and biofilm cells at different growth stages enabled a robust response function without further optimization. During stationary phase, a low and slow response was observed. Interestingly, the IPTG and Cuma sensors induced a higher stationary phase response in ASW cultures compared to those in rich medium. This may be due to lower oxygen tension in rich medium leading to low overall protein expression, while the lower cell numbers in ASW allowed for protein expression even when cells were growth limited. Sensors were also activated in *M. atlanticus* during early stages of biofilm formation on carbon electrodes in seawater medium, consistent with previous results using the IPTG sensor in an electrochemical flow cell configuration ^15^.

Although expression of YFP was relatively straightforward, control of electrical current production using the same sensors was more complex. Previous *in vitro* work has suggested that CymA uses MK-7 as a cofactor to accept electrons from the quinone pool ^34^ (Figure 2B). Additionally, studies in *S. oneidensis* and in *E. coli* have shown that mutants defective in the menaquinone synthesis pathway were defective in current production ^11, 36^. Results presented here show for the first time that the *So*EET pathway can be functionally expressed in an organism that naturally does not produce MK. However, exogenously added MK-4 was required for significantly higher current production. It might be possible to express the MK biosynthesis pathway in *M. atlanticus*, and remove the requirement for exogenous MK. It was recently shown that the entire MK biosynthesis pathway from *E. coli* could be ported to *Rhodobacter sphaeroides* to produce MK ^40^.

Engineered *M. atlanticus* biofilms use direct electron transfer, rather than small molecule mediators, to generate current. This is suggested by the rapid current recovery seen after medium replacement in Figure 6F, in contrast to a loss of current seen in medium replacement experiments in *S. oneidensis* biofilms, where soluble shuttles appear to play a large role in overall current production ^31^. The relative imperviousness of the current to medium exchanges may prove advantageous in sensor designs where the biofilm electrode is moved between different samples, or is configured as a remote sensor where the sampling matrix is continuously refreshed. *M. atlanticus* biofilms self-assemble and preinduction of *So*EET proteins is not required for increased current. This is an important step in the search for biosensors: they should ideally be able to function in environmental sample matrices. In contrast, *E. coli* engineered to express the Mtr and CymA proteins have used cells grown aerobically on rich medium and preinduced to measure the increase in current ^9, 41^ or additional engineering to immobilize cells at the electrode ^42^. Inducible current during biofilm formation has been demonstrated in *S. oneidensis* with a refactored native promoter for trimethylamine N-oxide (TMAO) ^6^, with the DAPG sensor used here ^16^, and with sensors for IPTG, Tc, and arabinose ^7^. However, current induction after biofilm formation was not assessed. Work in *E. coli* has focused primarily on increasing the efficiency of electricity production ^43^, rather than focusing on the ability to tune and temporally control the current produced. In all cases sensing after biofilm formation is slow and inconsistent, but recent progress on creating real-time response circuits is promising as a way to actuate current on demand ^12^.

CV of the CP1-eet strain shows a catalytic wave with an E_M_ of -0.081 V, which suggests that the MtrCAB complex is embedded in the cell’s outer membrane and can transfer electrons directly to the electrode. Our observed E_M_ is in range of that reported for purified MtrC ^29, 44^, and the MtrCAB complex in proteoliposomes ^30^. *E. coli* engineered to express the MtrCAB conduit and CymA showed a catalytic wave centered near 0.210 V SHE (0 V Ag/AgCl) ^10^. More recently, a redox potential of 0.051 V SHE was reported using differential pulse voltammetry (DPV) for an improved strain of *E. coli* expressing a modified version of the *ccm* operon ^41^.

Differences in the reported redox potential could be due to any number of factors that influence the equilibrium potential of the MtrCAB conduit when embedded in the cell membrane. For example, CV of *S. oneidensis* whole cells typically shows multiple redox features due to flavin cofactors and additional *c-*type cytochromes that bind to the MtrC conduit ^45-46^, making it difficult to compare to our data. While MK added to the bioelectrochemical reactors results in a pronounced set of reversible electrochemical peaks, attributed to direct interaction of MK with the electrode, a catalytic wave is only observed for induced CP1-eet. Moreover, when MK was added to high cell density cultures prior to inoculation of the bioelectrochemical reactors, there was little to no electrochemical signal attributable to MK, presumably because it partitioned into the cells, leaving a clearly defined catalytic wave attributed here to induced SoEET. Finally, electron transfer through the *So*EET conduit was reversible where CV indicated a shift in the onset potential for O_2_ reduction. Reversibility of the *So*EET pathway has also been shown in both the native host ^38-39^ and in *E. coli* engineered with the *So*EET pathway ^11^. The ability to engineer electron uptake into cells is important because it enables bidirectional communication with cells, where a response signal generated by a machine could direct cells on what to do next ^47^. The reverse electron flow did not require addition of MK, suggesting EET can occur from CymA to the endogenous quinone pool (ubiquone) and consistent with reliance on exogenous MK to enable EET in the forward direction due to its lower formal potential. This type of direct communication with biofilms would be in contrast to electrogenetic systems ^48^, in which electronic communication is achieved with planktonic cells via a soluble mediator. Delivery of excess reducing equivalents to cells also has important implications for microbial electrosynthesis ^49^ and electrofermentation ^50^. Although wild type *M. atlanticus* biofilms have been shown to take up electrons for O_2_ reduction when configured as a cathode ^17^, more work is needed to understand where electrons go and how this could be harnessed using synthetic biology.

## Conclusion

Engineered electroactive bacteria are envisioned for use as living sensors for the marine environment. Here we demonstrated the modularity of genetically encoded sensors optimized for *E. coli* in the marine chassis *M. atlanticus*. Sensors were used to show that the *S. oneidensis* electron transfer conduit could be actuated under operationally relevant conditions, a key step toward application. In addition to the *S. oneidensis* EET proteins, overexpression of the cytochrome *c* maturation pathway was required for current production in *M. atlanticus*, confirming design requirements previously realized in *E. coli*. Addition of exogenous menaquinone was also required to maximize current and future work should address whether this need can be fulfilled genetically through additional strain engineering.

## Materials and Methods

### Media and growth conditions

*M. atlanticus* CP1 was grown on BB medium or artificial seawater (ASW) as specified. Unless otherwise noted, chemicals came from Millipore-Sigma (St. Louis, Mo). BB medium consisted of 18.5 g Difco Marine (BD Biosciences), 2.5 g yeast extract, 5 g sodium chloride, and 5 g tryptone per liter. For solid medium, 1.5% agar added. Artificial seawater (ASW) medium contained 27.50 g NaCl, 3.80 g MgCl_2_•6H_2_O, 6.78 g MgSO_4_• H_2_O, 0.72 g KCl, 0.62 g NaHCO_3_, 5.58 g CaCl_2_• H_2_O, 1.00 g NH_4_Cl, 0.05 g K_2_HPO_4_, 1 mL Wolfe’s Trace Mineral Solution per liter ^51^, and 50 mM lactate. The NaHCO_3_ was autoclaved in a separate sealed bottle and added after autoclaving and cooling to avoid precipitation. The final medium pH was between 6.5 and 6.8. For planktonic growth in ASW (during plate assay and flow cytometry), the calcium was reduced to 0.05 g/L, as the calcium concentration in the standard ASW medium encouraged biofilm formation rather than planktonic growth ^17^. When added, IPTG (Gold Biotechnology, St. Louis, MO) was added from stock solutions prepared in water, DAPG (Santa Cruz Biotechnology, Dallas, TX) was added from stock solutions prepared in dimethyl formamide, tetracycline was added from stock solutions prepared in 80% ethanol, p-cumate was added from stock solutions prepared in 80% ethanol, Menaquinone-4 was added from a 20 mM stock solution in dimethyl sulfoxide, and Riboflavin was added from a freshly prepared 100 µM stock solution in water.

### Molecular techniques

Plasmids and strains used in this study are listed in Table 1, and primers used are shown in Supplemental Table 1.

All fragments for cloning were amplified using Q5 polymerase 2X master mix (New England Biolabs, Ipswich, Ma), and run on a 1% agarose gel with ethidium bromide. The band was cut out of the gel and purified using the Wizard gel purification kit (Promega). Each purified fragment was cloned into the amplified and purified backbone with Gibson assembly ^52^ using reagents from New England Biolabs, and transformed into *E. coli* DH5α via electroporation. The resulting plasmid was checked with colony PCR and Sanger sequencing. Colony PCR was performed by boiling a single colony in 10 mM NaOH, and using 1 µL of the supernatant in a 50 µL PCR reaction. Sanger sequencing was performed by Eurofins or Genewiz. Correct plasmids were purified using a miniprep kit (Qiagen) and chemically transformed into *E. coli* WM3064, then conjugated into *M. atlanticus* strains as previously described ^47^. The resulting transformants were restruck on antibiotic plates twice to obtain a pure strain.

Two different background strains were used to assess promoter response for the sensors: wild type *M. atlanticus* was used in planktonic culture experiments (plate reader and flow cytometry assays), and a strain harboring a genomic insert of the red fluorescent protein dTomato ^17^ was used for microscopy experiments. Each sensor cassette was cloned into the broad range vector pBBr1-mcs2, as shown in Figure 1A. The *ccm* operon was initially cloned into the pSEVA351 plasmid ^26^, which carries chloramphenicol resistance. However, we determined that the presence of chloramphenicol in the reactor caused a significant current increase even in controls containing pSEVA351 with *yfp* instead of the *ccm* operon (Supplemental Figure 8A). We therefore switched the chloramphenicol resistance in the plasmid to gentamycin, as gentamycin does not have an effect on current production (Supplemental Figure 8B).

Chromosomal insertions were made in the intergenic region downstream of the *glm* operon (position 1926204 in the genome) as described previously ^17^. We checked this insertion site for read-through from the upstream promoter by constructing a strain with *yfp* controlled by the TetR promoter inserted at this position, and confirming that fluorescence was not observed in the absence of aTc (Supplemental Figure 9). Steps for performing chromosomal insertions were as described above, with the following alterations: the plasmid backbone used for the insertions was pJQ200sk, which does not replicate in *M. atlanticus* and has the *gent*^*R*^ and *sacB* genes. After conjugation and restreaking on selective plates, a single colony of the transformed strain was grown for two days in BB broth without gentamycin. The culture was spread on BB plates with 10% sucrose and incubated for 2 days at 30°C. Colonies were restruck on BB plates, and tested for successful insertion with colony PCR of the insertion region.

The *M. atlanticus mtr* strain was made as described above by using the genes encoding the outer membrane metal-reducing (*mtr*) operon ^20-21^.

### Fluorescence measurements

To quantify YFP fluorescence in response to each inducer, strains transformed with the relevant cassette were inoculated from a BB-medium plate into a 5 mL liquid culture (BB or ASW + 50 mM lactate medium) and grown at 30°C overnight. The next day, the cultures were diluted 1:50 into the appropriate medium and aliquoted into 500 µL aliquots in a sterile 48-well plate (Corning, Corning, NY). The sides of the plate were sealed with lab tape (VWR) to prevent excessive evaporation over the course of the experiment. The plate was incubated with shaking in a plate reader at 30°C, with absorbance and YFP fluorescence measured at 10 minute intervals. After 6 hours of growth in the plate reader, the appropriate amount of inducer was added to each well, and the plate was returned to the plate reader. To make the titrations shown in Figure 1B, the concentration of inducer was plotted on the x-axis versus the measured fluorescence 21 hours after inoculation, once the fluorescence level had stabilized. The curves were fit using excel with the Solver add-in according to the methods in Meyer et al ^19^. The values for K, ymin, ymax, and n are reported in Supplemental Table 2.

Flow cytometry was performed as follows; cultures were grown overnight at 30°C in tubes and diluted 1:50 into a 0.5 mL volume in a 48-well plate with the inducer added immediately. After a 7-hour induction for BB broth and 18 hours for ASW, the cultures were run on a flow cytometer (Accuri c6 plus with autosampler, BD Biosciences). The time of measurement was chosen for each medium type to provide enough cells for measurement while avoiding clumping and surface attachment, which *M. atalanticus* is prone to.

To test induction during stationary phase, cultures were inoculated as described above, but were incubated for 18 hours before induction in order to induce at early stationary phase. Fluorescence measurements from 14 hours after induction were plotted and fitted as described above.

### BES setup

The BES were run in 170 mL electrochemical reactors (Pine Instruments, Grove City PA). The working and counter electrodes were made of 20 cm^2^ acid washed carbon cloth attached to titanium wire via titanium screws and nuts. The reference electrodes were Ag/AgCl, and potentials are reported here as vs. SHE by adding 0.21V to the experimental potentials. Reactors were held at 30°C with water jackets, and stirred with magnetic stir bars. The setup is shown in Supplemental Figure 4. Potential was maintained and current output recorded with a multichannel VMP3 potentiostat (Biologic, Seyssinet-Pariset, France). The inducer molecules were added from 10,000x stock solutions and the antibiotics were added from 1000x stock solutions. MK was added from a 10,000x stock solution in DMSO to a final concentration of 2 µM. The reactors were tested by performing cyclic voltammetry at 10 mV/S from 0.21 to 0.51 V SHE to check response of electrodes before inoculating. The reactors were held at 0.51 V for 30 minutes to allow current to stabilize, then inoculated. The first hour of current (encompassing initial startup and inoculation) was not plotted, to remove initial noise. Rubber stoppers (VWR) were used for all the reactor openings, restricting air exchange. The inoculated reactors were allowed to grow for 50 hours, during which time they became visibly cloudy, and current production was recorded. After 50 hours, a biotic CV was run at 0.2 mV/S, to characterize the biofilms.

When rubber stoppers were used multiple times, inconsistent current was observed, likely due to a loosening and cracking of the puncture through which the wire was inserted for the counter and working electrode. Previous experiments with wild type *M. atlanticus* indicated that current production only begins when the oxygen tension in the reactor drops ^14^, and that the number times the stopper was used affected the oxygen concentration (Elizabeth Onderko, unpublished observation). We therefore only reused punctured stoppers twice before replacing with new ones for the working and counter electrodes.

### Microscopy

For fluorescence visualization on electrodes, *M. atlanticus* CP-1 with dTomato expressed constitutively from a chromosomal insertion and the sensor carried on the plasmid were inoculated into reactors set up and run as described above. The appropriate inducer molecule was added to the reactor at the time of inoculation. After 24 hours, the carbon cloth was removed and a piece cut with a sterile razor blade for imaging. The sample was placed in a welled sample slide, and imaged on a Zeiss Axio Observer.Z1/7 with the Plan-Apochromat 40X oil objective. The laser wavelengths were 488 nm at 0.55% power for YFP detection with a detection wavelength of 464-592, and 561 nm at 0.2% power for dTomato with a detection range of 592-700. The exposure setting for each fluorophore was maintained across all samples. Images shown were representative of three images taken per sample, and two reactors were run for each condition tested.

### Heme stain

Cell pellets were spun down from 5 mL cultures and resuspended in 0.5 mL 100 mM Tris buffer, pH7. The resuspended pellets were sonicated for 30 seconds in 3 second bursts with 2 second rests on ice. NuPage 4X sample buffer (ThermoFisher, Waltham, MA) without reducing agent was added to the lysate, and incubated at 95°C for 10 minutes, then loaded on a NuPage 4-12% protein gel. The amount loaded per well was normalized to 30 µL (sample and buffer) of lysate with 10 mg/mL protein, measured with 280 unadjusted absorbance on a Nanodrop. The gel was then rinsed briefly in water, soaked in 12.5% tri-chloroacetic acid for 30 minutes, and rinsed in deionized water for 30 minutes. The heme stain was performed by adding 200 mg *o*-dianisidine to 180 mL diH2O, and stirring rapidly for 30 minutes. 20 mL of 0.5 M Na-citrate buffer, pH 4.4 was added, followed by 1 mL of 30% H_2_O_2_. The mixture was immediately poured over the gel. The gel was incubated in a covered container with shaking at 25°C and checked periodically. Gels were imaged after 1 hour.

### Protein Identification by Liquid Chromatography Tandem Mass Spectrometry (LC-MS/MS)

Bacterial strains were cultured in 5 mL BB broth and inducers overnight with shaking at 30°C. Cultures were rinsed, pelleted, and frozen at -80°C prior to sample processing via Pressure Cycling Technology (PCT), as previously described ^53^. Frozen samples were thawed and reconstituted in a solvent of 40 mM ammonium bicarbonate, 10% n-propanol and lysed via barocycler (Pressure BioSciences, South Easton, MA). For lysis, 60 cycles of PCT were conducted at 25°C, 45 kpsi for 25 seconds, with 10 second pauses between cycles. Extracted protein samples were reduced and alkylated with 10 mM DTT and 30 mM iodoacetamide, respectively, and incubated in the dark at ambient laboratory temperature for 30 minutes. For proteolytic digestion with trypsin, 60 cycles of PCT were conducted at 45°C, 45 kpsi for 50 seconds, with 10-second pauses between cycles. Liquid chromatography, tandem mass spectrometry (LC-MS/MS) was performed on a Orbitrap Fusion Lumos Tribrid mass spectrometer coupled to an Ultimate 3000 nanoRSLC system (Thermo Scientific, Waltham, MA, USA), as previously described ^54-55^. Mass spectrometry data were transformed and raw data extracted into peak lists via in-house scripting ^56^. Protein identification assignment to the predicted *M. atlanticus* proteome ^57^ was performed using a combination of Mascot Server (version 2.6.2, Matrix Science, London, UK) and ScaffoldQ+S (v. 4.8.9, Proteome Software, Portland, OR). The *M. atlanticus* protein database searched was retrieved on 04 JAN 2021, and its 4,107 entries were concatenated to 8 recombinant, plasmid-encoded proteins, and a manually curated inventory of 190 common contaminant proteins (keratins, trypsin, etc.) ^56^.

Raw proteomics data was deposited in the Mass Spectrometry Interactive Virtual Environment (MassIVE) repository with the accession number MSV000090010.

## Acknowledgements

This work was funded through NRL’s 6.1 base program and the Applied Research for the Advancement of Science and Technology Priorities Program on Synthetic Biology for Military Environments.

## Citations

1. Bond, D. R.; Holmes, D. E.; Tender, L. M.; Lovley, D. R., Electrode-reducing microorganisms that harvest energy from marine sediments. Science 2002, 295 (5554), 483–485.

2. Tender, L. M.; Reimers, C. E.; Stecher, H. A.; Holmes, D. E.; Bond, D. R.; Lowy, D. A.; Pilobello, K.; Fertig, S. J.; Lovley, D. R., Harnessing microbially generated power on the seafloor. Nat Biotechnol 2002, 20 (8), 821–825.

3. Tender, L. M.; Gray, S. A.; Groveman, E.; Lowy, D. A.; Kauffman, P.; Melhado, J.; Tyce, R. C.; Flynn, D.; Petrecca, R.; Dobarro, J., The first demonstration of a microbial fuel cell as a viable power supply: Powering a meteorological buoy. J Power Sources 2008, 179 (2), 571–575.

4. Eddie, B. J.; Wang, Z.; Malanoski, A. P.; Hall, R. J.; Oh, S. D.; Heiner, C.; Lin, B.; Strycharz-Glaven, S. M., ‘Candidatus Tenderia electrophaga’, an uncultivated electroautotroph from a biocathode enrichment. Int J Syst Evol Microbiol 2016, 66 (6), 2178–2185.

5. Eddie, B. J.; Wang, Z.; Hervey, W. J. t.; Leary, D. H.; Malanoski, A. P.; Tender, L. M.; Lin, B.; Strycharz-Glaven, S. M., Metatranscriptomics Supports the Mechanism for Biocathode Electroautotrophy by “Candidatus Tenderia electrophaga”. mSystems 2017, 2 (2).

6. West, E. A.; Jain, A.; Gralnick, J. A., Engineering a Native Inducible Expression System in Shewanella oneidensis to Control Extracellular Electron Transfer. Acs Synth Biol 2017, 6 (9), 1627–1634.

7. Cao, Y.; Song, M.; Li, F.; Li, C.; Lin, X.; Chen, Y.; Chen, Y.; Xu, J.; Ding, Q.; Song, H., A Synthetic Plasmid Toolkit for Shewanella oneidensis MR-1. Front Microbiol 2019, 10, 410.

8. Goldbeck, C. P.; Jensen, H. M.; TerAvest, M. A.; Beedle, N.; Appling, Y.; Hepler, M.; Cambray, G.; Mutalik, V.; Angenent, L. T.; Ajo-Franklin, C. M., Tuning Promoter Strengths for Improved Synthesis and Function of Electron Conduits in Escherichia coli. Acs Synth Biol 2013, 2 (3), 150–159.

9. TerAvest, M. A.; Zajdel, T. J.; Ajo-Franklin, C. M., The Mtr Pathway of Shewanella oneidensis MR-1 Couples Substrate Utilization to Current Production in Escherichia coli. Chemelectrochem 2014, 1 (11), 1874–1879.

10. Jensen, H. M.; TerAvest, M. A.; Kokish, M. G.; Ajo-Franklin, C. M., CymA and Exogenous Flavins Improve Extracellular Electron Transfer and Couple It to Cell Growth in Mtr-Expressing Escherichia coli. Acs Synth Biol 2016, 5 (7), 679–688.

11. Baruch, M.; Tejedor-Sanz, S.; Su, L.; Ajo-Franklin, C. M., Electronic control of redox reactions inside Escherichia coli using a genetic module. PLoS One 2021, 16 (11), e0258380.

12. Atkinson, J. T.; Su, L.; Zhang, X.; Bennett, G. N.; Silberg, J. J.; Ajo-Franklin, C. M., Real-time environmental monitoring of contaminants using living electronic sensors. bioRxiv 2021, 2021.06.04.447163.

13. Wang, Z.; Leary, D. H.; Malanoski, A. P.; Li, R. W.; Hervey, W. J. t.; Eddie, B. J.; Tender, G. S.; Yanosky, S. G.; Vora, G. J.; Tender, L. M.; Lin, B.; Strycharz-Glaven, S. M., A previously uncharacterized, nonphotosynthetic member of the Chromatiaceae is the primary CO2-fixing constituent in a self-regenerating biocathode. Appl Environ Microb 2015, 81 (2), 699–712.

14. Onderko, E. L.; Phillips, D. A.; Eddie, B. J.; Yates, M. D.; Wang, Z.; Tender, L. M.; Glaven, S. M., Electrochemical Characterization of Marinobacter atlanticus Strain CP1 Suggests a Role for Trace Minerals in Electrogenic Activity. Frontiers in Energy Research 2019, 7.

15. Phillips, D. A.; Bird, L. J.; Eddie, B. J.; Yates, M. D.; Tender, L. M.; Voigt, C. A.; Glaven, S. M., Activation of Protein Expression in Electroactive Biofilms. Acs Synth Biol 2020, 9 (8), 1958–1967.

16. Yates, M. D.; Bird, L. J.; Eddie, B. J.; Onderko, E. L.; Voigt, C. A.; Glaven, S. M., Nanoliter scale electrochemistry of natural and engineered electroactive bacteria. Bioelectrochemistry 2021, 137, 107644.

17. Bird, L. J.; Wang, Z.; Malanoski, A. P.; Onderko, E. L.; Johnson, B. J.; Moore, M. H.; Phillips, D. A.; Chu, B. J.; Doyle, J. F.; Eddie, B. J.; Glaven, S. M., Development of a Genetic System for Marinobacter atlanticus CP1 (sp. nov.), a Wax Ester Producing Strain Isolated From an Autotrophic Biocathode. Frontiers in Microbiology 2018, 9.

18. Bird, L. J.; Onderko, E. L.; Phillips, D. A.; Mickol, R. L.; Malanoski, A. P.; Yates, M. D.; Eddie, B. J.; Glaven, S. M., Engineered living conductive biofilms as functional materials. Mrs Commun 2019, 9 (2), 505–517.

19. Meyer, A. J.; Segall-Shapiro, T. H.; Glassey, E.; Zhang, J.; Voigt, C. A., Escherichia coli “Marionette” strains with 12 highly optimized small-molecule sensors. Nat Chem Biol 2019, 15 (2), 196–204.

20. Richardson, D. J.; Butt, J. N.; Fredrickson, J. K.; Zachara, J. M.; Shi, L.; Edwards, M. J.; White, G.; Baiden, N.; Gates, A. J.; Marritt, S. J.; Others, The ‘porin--cytochrome’model for microbe-to-mineral electron transfer. Mol. Microbiol. 2012, 85 (2), 201–212.

21. Edwards, M. J.; White, G. F.; Lockwood, C. W.; Lawes, M. C.; Martel, A.; Harris, G.; Scott, D. J.; Richardson, D. J.; Butt, J. N.; Clarke, T. A., Structural modeling of an outer membrane electron conduit from a metal-reducing bacterium suggests electron transfer via periplasmic redox partners. J. Biol. Chem. 2018, 293 (21), 8103–8112.

22. Sturm, G.; Richter, K.; Doetsch, A.; Heide, H.; Louro, R. O.; Gescher, J., A dynamic periplasmic electron transfer network enables respiratory flexibility beyond a thermodynamic regulatory regime. ISME J. 2015, 9 (8), 1802–1811.

23. Schwalb, C.; Chapman, S. K.; Reid, G. A., The tetraheme cytochrome CymA is required for anaerobic respiration with dimethyl sulfoxide and nitrite in Shewanella oneidensis. Biochemistry 2003, 42 (31), 9491–9497.

24. Marritt, S. J.; Lowe, T. G.; Bye, J.; McMillan, D. G.; Shi, L.; Fredrickson, J.; Zachara, J.; Richardson, D. J.; Cheesman, M. R.; Jeuken, L. J.; Butt, J. N., A functional description of CymA, an electron-transfer hub supporting anaerobic respiratory flexibility in Shewanella. Biochem J 2012, 444 (3), 465–74.

25. Jensen, H. M.; Albers, A. E.; Malley, K. R.; Londer, Y. Y.; Cohen, B. E.; Helms, B. A.; Weigele, P.; Groves, J. T.; Ajo-Franklin, C. M., Engineering of a synthetic electron conduit in living cells. Proceedings of the National Academy of Sciences of the United States of America 2010, 107 (45), 19213–19218.

26. Silva-Rocha, R.; Martinez-Garcia, E.; Calles, B.; Chavarria, M.; Arce-Rodriguez, A.; de Las Heras, A.; Paez-Espino, A. D.; Durante-Rodriguez, G.; Kim, J.; Nikel, P. I.; Platero, R.; de Lorenzo, V., The Standard European Vector Architecture (SEVA): a coherent platform for the analysis and deployment of complex prokaryotic phenotypes. Nucleic Acids Res 2013, 41 (Database issue), D666–75.

27. Strycharz, S. M.; Malanoski, A. P.; Snider, R. M.; Yi, H.; Lovley, D. R.; Tender, L. M., Application of cyclic voltammetry to investigate enhanced catalytic current generation by biofilm-modified anodes of Geobacter sulfurreducens strain DL1 vs. variant strain KN400. Energ Environ Sci 2011, 4 (3), 896–913.

28. Torres, C. I.; Marcus, A. K.; Parameswaran, P.; Rittmann, B. E., Kinetic experiments for evaluating the Nernst-Monod model for anode-respiring bacteria (ARB) in a biofilm anode. Environmental Science & Technology 2008, 42 (17), 6593–6597.

29. Firer-Sherwood, M.; Pulcu, G. S.; Elliott, S. J., Electrochemical interrogations of the Mtr cytochromes from Shewanella: opening a potential window. J Biol Inorg Chem 2008, 13 (6), 849–854.

30. Hartshorne, R. S.; Reardon, C. L.; Ross, D.; Nuester, J.; Clarke, T. A.; Gates, A. J.; Mills, P. C.; Fredrickson, J. K.; Zachara, J. M.; Shi, L.; Beliaev, A. S.; Marshall, M. J.; Tien, M.; Brantley, S.; Butt, J. N.; Richardson, D. J., Characterization of an electron conduit between bacteria and the extracellular environment. Proceedings of the National Academy of Sciences of the United States of America 2009, 106 (52), 22169–22174.

31. Kotloski, N. J.; Gralnick, J. A., Flavin electron shuttles dominate extracellular electron transfer by Shewanella oneidensis. Mbio 2013, 4 (1).

32. Madsen, C. S.; TerAvest, M. A., NADH dehydrogenases Nuo and Nqr1 contribute to extracellular electron transfer by Shewanella oneidensis MR-1 in bioelectrochemical systems. Sci Rep-Uk 2019, 9 (1), 14959.

33. Duhl, K. L.; Tefft, N. M.; TerAvest, M. A., Shewanella oneidensis MR-1 Utilizes both Sodium- and Proton-Pumping NADH Dehydrogenases during Aerobic Growth. Appl Environ Microb 2018, 84 (12).

34. McMillan, D. G.; Marritt, S. J.; Butt, J. N.; Jeuken, L. J., Menaquinone-7 is specific cofactor in tetraheme quinol dehydrogenase CymA. J Biol Chem 2012, 287 (17), 14215–25.

35. Urban, P. F.; Klingenberg, M., On Redox Potentials of Ubiquinone and Cytochrome B in Respiratory Chain. Eur J Biochem 1969, 9 (4), 519-+.

36. Mevers, E.; Su, L.; Pishchany, G.; Baruch, M.; Cornejo, J.; Hobert, E.; Dimise, E.; Ajo-Franklin, C. M.; Clardy, J., An elusive electron shuttle from a facultative anaerobe. Elife 2019, 8.

37. Dharmaraj, K.; Roman Silva, J. I.; Kahlert, H.; Lendeckel, U.; Scholz, F., The acid-base and redox properties of menaquinone MK-4, MK-7, and MK-9 (vitamin K2) in DMPC monolayers on mercury. Eur Biophys J 2020, 49 (3-4), 279–288.

38. Rowe, A. R.; Rajeev, P.; Jain, A.; Pirbadian, S.; Okamoto, A.; Gralnick, J. A.; El-Naggar, M. Y.; Nealson, K. H., Tracking Electron Uptake from a Cathode into Shewanella Cells: Implications for Energy Acquisition from Solid-Substrate Electron Donors. Mbio 2018, 9 (1).

39. Coursolle, D.; Gralnick, J. A., Modularity of the Mtr respiratory pathway of Shewanella oneidensis strain MR-1. Mol Microbiol 2010, 77 (4), 995–1008.

40. Jun, D.; Richardson-Sanchez, T.; Mahey, A.; Murphy, M. E. P.; Fernandez, R. C.; Beatty, J. T., Introduction of the Menaquinone Biosynthetic Pathway into Rhodobacter sphaeroides and de Novo Synthesis of Menaquinone for Incorporation into Heterologously Expressed Integral Membrane Proteins. Acs Synth Biol 2020, 9 (5), 1190–1200.

41. Su, L.; Fukushima, T.; Ajo-Franklin, C. M., A hybrid cyt c maturation system enhances the bioelectrical performance of engineered Escherichia coli by improving the rate-limiting step. Biosensors & Bioelectronics 2020, 165.

42. Lienemann, M.; TerAvest, M. A.; Pitkanen, J. P.; Stuns, I.; Penttila, M.; Ajo-Franklin, C. M.; Jantti, J., Towards patterned bioelectronics: facilitated immobilization of exoelectrogenic Escherichia coli with heterologous pili. Microbial Biotechnology 2018, 11 (6), 1184–1194.

43. Li, F.; Li, Y.; Sun, L.; Chen, X.; An, X.; Yin, C.; Cao, Y.; Wu, H.; Song, H., Modular Engineering Intracellular NADH Regeneration Boosts Extracellular Electron Transfer of Shewanella oneidensis MR-1. ACS Synth. Biol. 2018, 7 (3), 885–895.

44. Hartshorne, R. S.; Jepson, B. N.; Clarke, T. A.; Field, S. J.; Fredrickson, J.; Zachara, J.; Shi, L.; Butt, J. N.; Richardson, D. J., Characterization of Shewanella oneidensis MtrC: a cell-surface decaheme cytochrome involved in respiratory electron transport to extracellular electron acceptors. J Biol Inorg Chem 2007, 12 (7), 1083–1094.

45. Okamoto, A.; Hashimoto, K.; Nealson, K. H.; Nakamura, R., Rate enhancement of bacterial extracellular electron transport involves bound flavin semiquinones. Proceedings of the National Academy of Sciences of the United States of America 2013, 110 (19), 7856–7861.

46. Nakamura, R.; Ishii, K.; Hashimoto, K., Electronic Absorption Spectra and Redox Properties of C Type Cytochromes in Living Microbes. Angewandte Chemie-International Edition 2009, 48 (9), 1606–1608.

47. Bird, L. J.; Kundu, B. B.; Tschirhart, T.; Corts, A. D.; Su, L.; Gralnick, J. A.; Ajo-Franklin, C. M.; Glaven, S. M., Engineering Wired Life: Synthetic Biology for Electroactive Bacteria. Acs Synth Biol 2021, 10 (11), 2808–2823.

48. Tschirhart, T.; Kim, E.; McKay, R.; Ueda, H.; Wu, H.-C.; Pottash, A. E.; Zargar, A.; Negrete, A.; Shiloach, J.; Payne, G. F.; Bentley, W. E., Electronic control of gene expression and cell behaviour in Escherichia coli through redox signalling. Nat. Commun. 2017, 8 (1), 14030.

49. Jourdin, L.; Burdyny, T., Microbial Electrosynthesis: Where Do We Go from Here? Trends Biotechnol 2021, 39 (4), 359–369.

50. Virdis, B.; D. Hoelzle R.; Marchetti, A.; Boto, S. T.; Rosenbaum, M. A.; Blasco-Gómez, R.; Puig, S.; Freguia, S.; Villano, M., Electro-fermentation: Sustainable bioproductions steered by electricity. Biotechnology Advances 2022, 59, 107950.

51. Emerson, D.; Floyd, M. M., Enrichment and isolation of iron-oxidizing bacteria at neutral pH. Methods Enzymol 2005, 397, 112–23.

52. Gibson, D. G.; Young, L.; Chuang, R. Y.; Venter, J. C.; Hutchison, C. A., 3rd; Smith, H. O., Enzymatic assembly of DNA molecules up to several hundred kilobases. Nat Methods 2009, 6 (5), 343–5.

53. Schultzhaus, J. N.; Dean, S. N.; Leary, D. H.; Hervey, W. J.; Fears, K. P.; Wahl, K. J.; Spillmann, C. M., Pressure cycling technology for challenging proteomic sample processing: application to barnacle adhesive. Integr Biol-Uk 2019, 11 (5), 235–247.

54. Dean, S. N.; Rimmer, M. A.; Turner, K. B.; Phillips, D. A.; Caruana, J. C.; Hervey, W. J.; Leary, D. H.; Walper, S. A., Lactobacillus acidophilus Membrane Vesicles as a Vehicle of Bacteriocin Delivery. Frontiers in Microbiology 2020, 11.

55. Schultzhaus, J. N.; Hervey, W. J.; Taitt, C. R.; So, C. R.; Leary, D. H.; Wahl, K. J.; Spillmann, C. M., Comparative analysis of stalked and acorn barnacle adhesive proteomes. Open Biol 2021, 11 (8).

56. Hervey, W. J.; Khalsa-Moyers, G.; Lankford, P. K.; Owens, E. T.; McKeown, C. K.; Lu, T. Y.; Foote, L. J.; Asano, K. G.; Morrell-Falvey, J. L.; McDonald, W. H.; Pelletier, D. A.; Hurst, G. B., Evaluation of Affinity-Tagged Protein Expression Strategies Using Local and Global Isotope Ratio Measurements. J Proteome Res 2009, 8 (7), 3675–3688.

57. Wang, Z.; Eddie, B. J.; Malanoski, A. P.; Hervey, W. J. t.; Lin, B.; Strycharz-Glaven, S. M., Complete Genome Sequence of Marinobacter sp. CP1, Isolated from a Self-Regenerating Biocathode Biofilm. Genome Announc 2015, 3 (5).

58. Woodcock, D. M.; Crowther, P. J.; Doherty, J.; Jefferson, S.; DeCruz, E.; Noyer-Weidner, M.; Smith, S. S.; Michael, M. Z.; Graham, M. W., Quantitative evaluation of Escherichia coli host strains for tolerance to cytosine methylation in plasmid and phage recombinants. Nucleic Acids Research 1989, 17 (9), 3469–3478.

59. Saltikov, C. W.; Newman, D. K., Genetic identification of a respiratory arsenate reductase. P Natl Acad Sci USA 2003, 100 (19), 10983–8.

60. Kovach, M. E.; Elzer, P. H.; Steven Hill, D.; Robertson, G. T.; Farris, M. A.; Roop, R. M.; Peterson, K. M., Four new derivatives of the broad-host-range cloning vector pBBR1MCS, carrying different antibiotic-resistance cassettes. Gene 1995, 166 (1), 175–176.

61. Quandt, J.; Hynes, M. F., Versatile suicide vectors which allow direct selection for gene replacement in gram-negative bacteria. Gene 1993, 127 (1), 15–21.

