## SUpplemental material for "A living conductive marine biofilm engineered to sense and respond to small molecules"

Naval Research Laboratory
4555 Overlook Ave.
Washington, DC 20375

**Supporting Information**

### Table of Contents

**Supplemental Table 1. Primers used in this study.**
**Supplemental Table 2: Induction parameters for promoters.**
**Supplemental Figure 1. Strain and sensor response to naringenin, vanillin, and choline.**
**Supplemental Figure 2. Response function in stationary phase in BB broth and ASW.**
**Supplemental Figure 3. Cytochrome *c* expression in CP1 strains.**
**Supplemental Figure 4. Schematic of electrochemical reactor.**
**Supplemental Figure 5. Heme stain of different strains after growth in BB broth.**
**Supplemental Figure 6. Abiotic MK pH dependency.**
**Supplemental Figure 7. CVs of high-density inoculum.**
**Supplemental Figure 8. Redox activity of Chloramphenicol.**
**Supplemental Figure 9. Inducible YFP inserted at the CP1 neutral site.**

**Supplemental Table 1: Primers used in this study**

| | | |
|---|---|---|
| pBBrF3 | acccgcgctcagccaacgatcgttaagcgggGCGTTAATATTTTGTTAAAATTCGC | cloning sensor cassette into pBBr |
| pBBrR2 | gggctcatgagcaaatattttatctgaggtAGCTGTTTCCTGTGTGAAATTG | cloning sensor cassette into pBBr |
| pAJMF2 | ttaacgcgaattttaacaaaatattaacgcCCCGCTTAACGATCGTTGGCTG | cloning sensor cassette into pBBr |
| pAJMR3 | agcggataacaatttcacacaggaaacagctACCTCAGATAAAATATTTGC | cloning sensor cassette into pBBr |
| CymAAJMF1 | taatactagagaaagaggggaaatactagcATGAACTGGCGTGCACTATT | CymA into DAPG sensor |
| pAJMcymAR1 | cgctgggtttaaatagtgcacgccagttcatGCTAGTATTTCCCCTCTTTCTC | CymA into DAPG sensor |
| CctACymAF | tcacccctatccaaaaggataaTACAATTGTGAGCAAAAAACTATTAAGTG | Connecting CymA to CctA |
| CymAMtr_R | tttccctgcataggtttggcaTTATCCTTTTGGATAGGGGTG | Connecting CymA to CctA |
| CctApbbjmR | cctcttttctggaatttggtaccGACTTGGCAATTACTTCTTCAG | cctA into DAPG sensor |
| pAJMCctA_F | ctgttctgaagaagtaattgCCAAGTCGGTACCAAATTCCAGAAAAG | cctA into DAPG sensor |
| pSEVA351 F | GTCGTGACTGGGAAAACCCTG | Sequencing pSEVA |
| pSEVA351R | GCCTCCTGTGTGAAATTGTTATC | Sequencing pSEVA |
| ptetRpSEVA351F | taaagcggataacaatttcacacaggaggcCCCGCTTAACGATCGTTGG | TetR cassette into pSEVA |
| tetRpSEVA351R | actagtcgccagggttttcccagtcacgacACCTCAGATAAAATATTTGCTCATG | TetR cassette into pSEVA |
| tetRCcmR1 | ataacggctcagacatCTAGTATTTCCCCTCTTTCTCTAG | ccm operon into TetR cassette |
| ccmtetRF1 | gaaagaggggaaatactagATGTCTGAGCCGTTATTACAG | ccm operon into TetR cassette |
| ccmtetRR1 | tctggaatttggtaccgagTCAGTCAATCAGTTGCTGATC | ccm operon into TetR cassette |
| tetRccmF1 | cgatcagcaactgattgactgaCTCGGTACCAAATTCCAG | ccm operon into TetR cassette |
| pSEVAchlorFWD | acccaagtaccgccacctaaTTTGACTTTTGTCCTTTTCCG | Switching pSEVA chloramphenicol resistance to gent |
| pSEVAhlorREV | ctgtacaaaaaaacagtcatTTTAGCTTCCTTAGCTCCTG | Switching pSEVA chloramphenicol resistance to gent |
| chlorpSEVAFWD | caggagctaaggaagctaaaATGACTGTTTTTTTGTACAGTCTATG | Switching pSEVA chloramphenicol resistance to gent |
| chlorpSEVAREV | ggaaaaggacaaaagtcaaaTTAGGTGGCGGTACTTGGGT | Switching pSEVA chloramphenicol resistance to gent |
| pSEVAchlorFWD | acccaagtaccgccacctaaTTTGACTTTTGTCCTTTTCCG | Switching pSEVA chloramphenicol resistance to gent |

**Supplemental Table 2**: **induction parameters for promoters.**

| **promoter** | **inducer** | **$y_{max}$ (AU)** | **$y_{min}$ (AU)** | **Dynamic range** | **K (µM)** | **n** |
|---|---|---|---|---|---|---|
| **Log phase** | | | | | | |
| $P_{tetR}$, BB broth | Tc | 12012 | 10 | 1201 | 1.7 | 3 |
| tetR, ASW | Tc | 2839 | 16 | 177 | 5.6 | 2.8 |
| lacI, BB broth | IPTG | 2960 | 10 | 296 | 44.7 | 2.2 |
| lacI, ASW | IPTG | 973 | 4 | 243 | 41.5 | 1.9 |
| cymR, BB broth | Cuma | 2490 | 6 | 407 | 1.4 | 2.4 |
| cymR ASW | Cuma | 1868 | 7 | 248 | 3.2 | 1.9 |
| phlF, BB broth | DAPG | 1685 | 3 | 527 | .75 | 1.7 |
| phlF, ASW | DAPG | 2013 | 7 | 288 | 1.7 | 2.8 |
| vanR, BB broth | Van | 1000 | 50 | 20 | 30 | 1.8 |
| ttgR, BB brohth | Nar | 45 | 260 | 0 | 45 | 2.5 |
| betI, BB broth | Cho | 300 | 200 | 2 | 6000 | 1.9 |
| **Stationary phase** | | | | | | |
| tetR, BB broth | Tc | 18 | 15 | 1 | 1.18 | 3 |
| tetR, ASW | Tc | 12 | 9 | 1 | 5.7 | .05 |
| lacI, BB broth | IPTG | 132 | 32 | 4 | 256.9 | 2.1 |
| lacI, ASW | IPTG | 481 | 8.4 | 57 | 38.1 | 1.4 |
| cymR, BB broth | Cuma | 260 | 17 | 15 | 6.8 | 2 |
| cymR ASW | Cuma | 1617 | 15 | 108 | 751 | 2.9 |
| phlF, BB broth | DAPG | 144 | 2 | 72 | 9.7 | 1.7 |
| phlF, ASW | DAPG | 75 | .06 | 1250 | 11.1 | 2 |

A

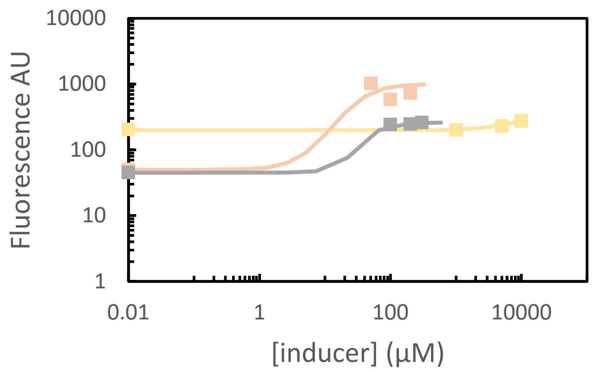


B

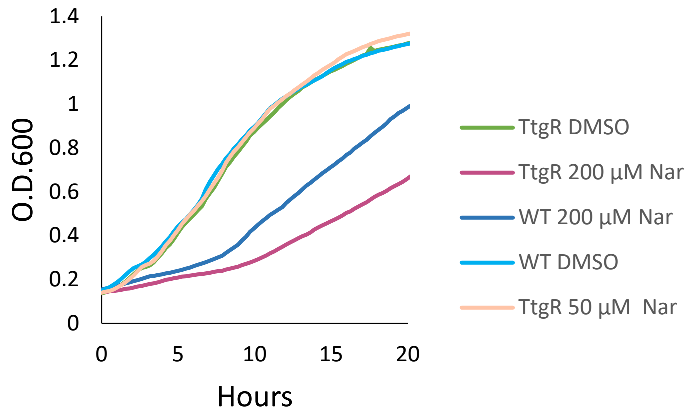


**Supplemental Figure 1. Strain and sensor response to naringenin (Nar), vanillin (Van) and choline (Cho).** A) Response function of naringenin (Nar), vanillic acid (Van), or choline chloride (Cho) sensor strains when exposed in rich medium at 20 hours: 0, 50, 100, and 200 µM Nar (peach); 0, 1,000, 5,000, and 10,000 µM Cho (yellow), 0, 100, 200, and 300 µM Van (grey). B) Growth curves (optical density at 600 nm (O.D.600) of the *M. atlanticus* wild type and naringenin (Nar) strain when exposed to dimethyl sulfoxide (DMSO) only or naringenin (50 or 200 µM) dissolved in DMSO.

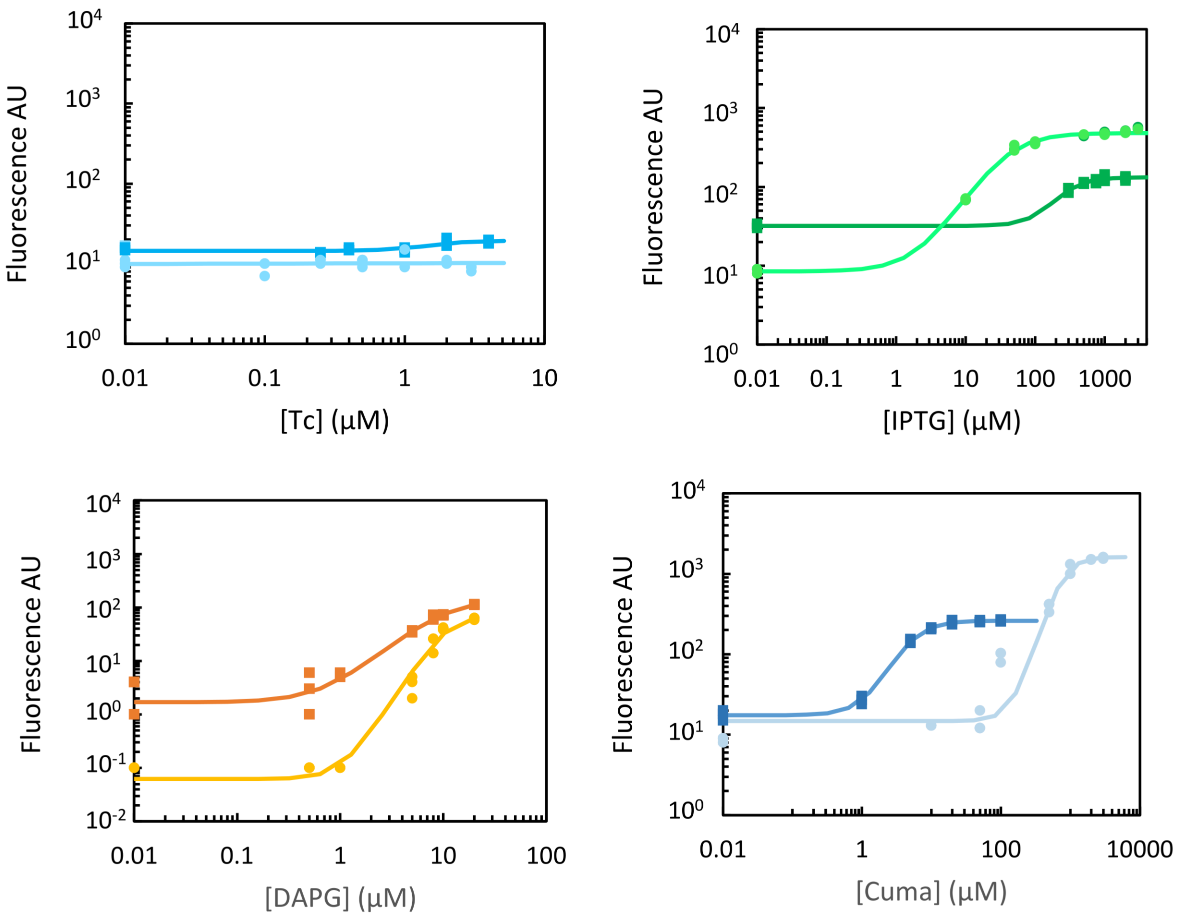


**Supplemental Figure 2. Response function in stationary phase in BB broth and ASW.** Inducer was added at 18 hours, during early stationary phase. The lighter color indicates induction in ASW, and the darker color indicates induction in BB broth. 3 biological replicates are shown individually.

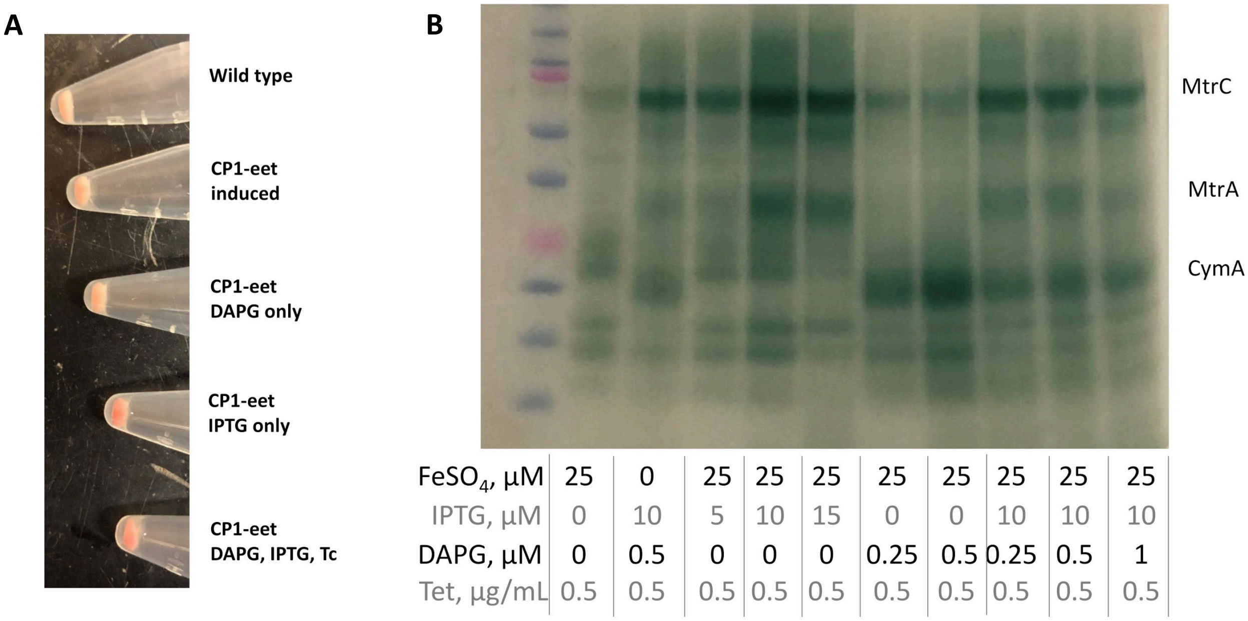


C

| | WT | CP1-eet uninduced | CP1-eet 0.5 µg/mL Tet | CP1-eet 0.25 µM DAPG 5 µM IPTG 0.5 µg/mL Tet | CP1-eet 0.5 µM DAPG 10 µM IPTG 0.5 µg/mL Tet |
|---|---|---|---|---|---|
| MtrC | 0 | 6.5 | 6.5 | 13.5 | 34.5 |
| CymA | 0 | 0 | 0 | 6 | 7 |
| MtrB | 0 | 3 | 3.5 | 4 | 13 |

**Supplemental Figure 3. Cytochrome *c* expression in CP1 strains.** A) Overnight expression of the Mtr proteins in BB broth makes the pellet appear redder. Inducer concentrations: DAPG = 0. 25 µM, IPTG = 10 µM, and Tc = 0.5 µg/mL. B) Expression of Mtr proteins and CymA is visible in overnight BB cultures. A slight increase in intensity is observed when comparing the same inductions conditions with and without additional iron added to the medium (lane 2 vs lane 9). Amount loaded per lane was normalized based on the 280 absorbance of cell lysates. C) Expression of MtrC, CymA, and MtrB proteins were identified by liquid chromatography tandem mass spectrometry (LC-MS/MS). Numbers indicate average hits from 2 biological replicates. Cultures were normalized by optical density.

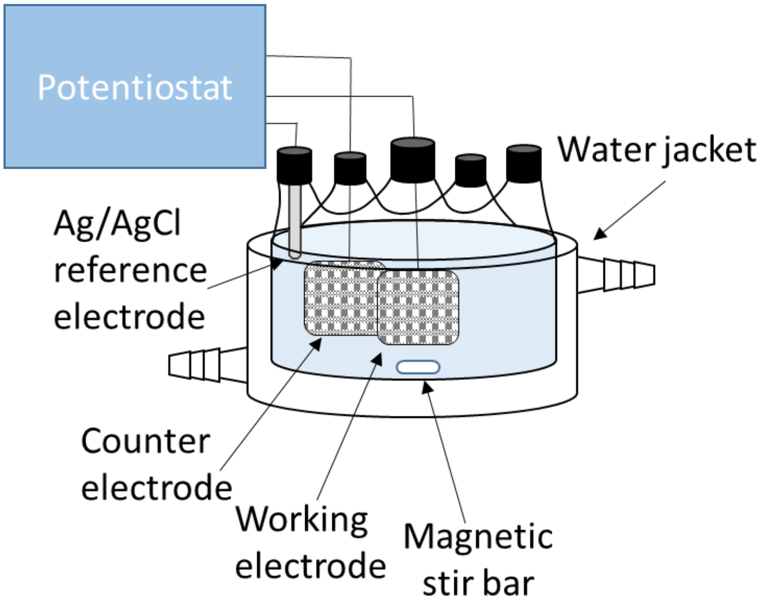


**Supplemental Figure 4**. **Schematic of electrochemical reactor**. Inner chamber is filled with 170 mL ASW medium, with the outer water jacket containing water at 30°C, continuously flowing from a heated water bath. Working and counter electrodes are made of carbon cloth and are controlled by a potentiostat.

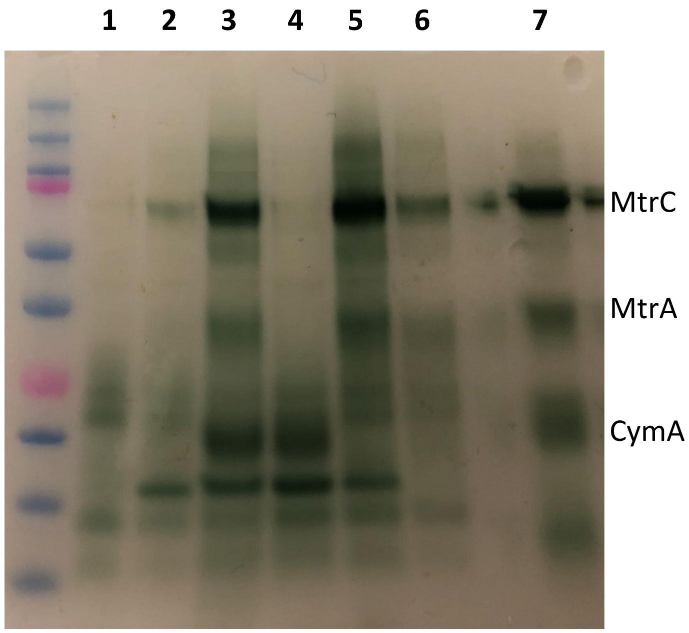


**Supplemental Figure 5. Heme stain of different strains after growth in BB broth with the concentrations used for electrode induction.** A) 1: *M. atlanticus* wild type; 2: CP1-eet uninduced; 3: CP1-eet induced; 4: CP1-cymAcctA induced; 5: CP1-mtr induced; 6: CP1-mcc induced; 7: Shewanella positive control. Induction = 10 µM IPTG, 0.25 µM DAPG, and 0.5 µg/mL Tc. The presence of the appropriate proteins in each strain indicates that the expression units (MtrCAB and CymA-CctA) are expressed correctly in the individual strains, even though no increase in current is observed.

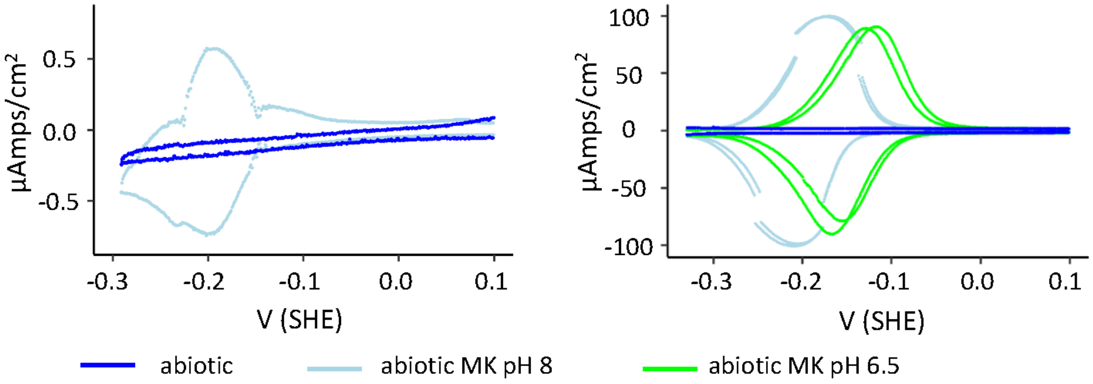


**Supplemental Figure 6. Abiotic MK pH dependency.** Top: CV of abiotic reactors with and without MK added. Bottom: Differential Pulse Voltammetry (DPV) of abiotic reactors with and without MK, and with the pH adjusted: after bubbling with nitrogen the medium in the abiotic reactor was pH 8. The pH was adjusted to 6.5, and the DPV with the adjusted pH is shown in green.

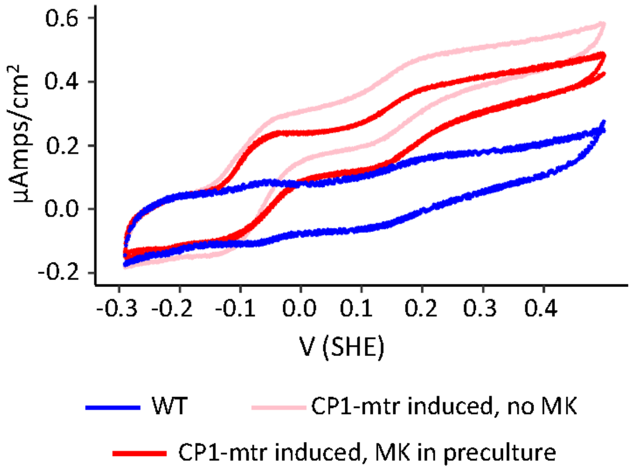


**Supplemental Figure 7. CVs of high-density inoculum.** 50 mL of preculture were grown and induced with DAPG, IPTG, Tc, and iron overnight. 2 hours before reactor inoculation, MK was added to the preculture. The culture was then pelleted, rinsed in ASW and resuspended in 5 ml ASW, and added to the reactor. Under these conditions, CP1-eet showed the catalytic wave at -0.09 V SHE both when MK was added to the preculture (red) and in a control to which MK was not added (pink), while wild type CP1 did not show the wave even when MK was added to the preculture (blue).

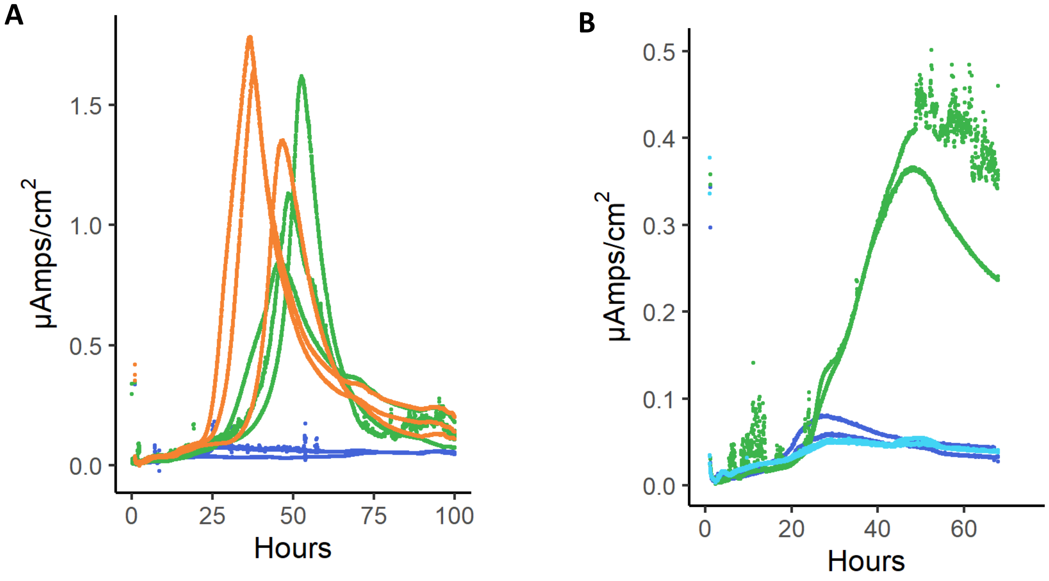


**Supplemental Figure 8. Redox activity of Chloramphenicol.** A) current produced with strains containing *so*EET proteins and Ccm proteins on a chloramphenicol resistant plasmid (orange) and a yfp containing control plasmid (green). Chloramphenicol boosts current production even in the absence of the EET proteins compared to wild type CP1 (blue). B) comparison of current production in resistant strain in the presence of gentamycin (light blue). Gentamycin does not cause the same current effects as chloramphenicol (green), but produced current similar to wt CP1 (blue).

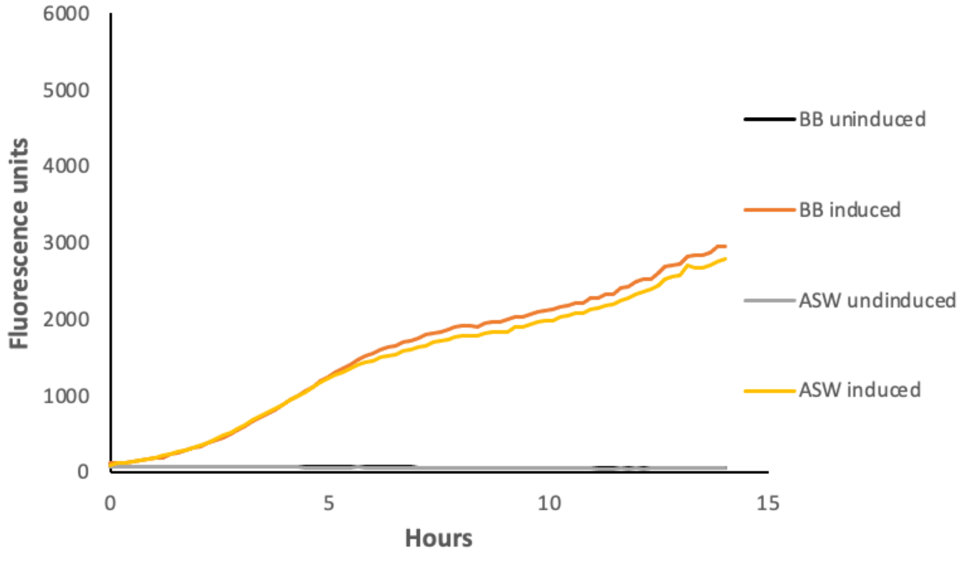


**Supplemental Figure 9. Inducible YFP inserted at the CP1 neutral site.** Chromosomal insertion of *yfp* under the Tc inducible promoter. The inducibility of the promoter when inserted in the chosen site indicates that there is no read through from nearby promoters on the chromosome.
